# Missense variant interaction scanning reveals a critical role of the FERM-F3 domain for tumor suppressor protein NF2 conformation and function

**DOI:** 10.1101/2022.12.11.519953

**Authors:** Christina S. Moesslacher, Jonathan Woodsmith, Elisabeth Auernig, Andreas Feichtner, Evelyne Jany-Luig, Stefanie Jehle, Josephine M. Worseck, Christian L. Heine, Eduard Stefan, Ulrich Stelzl

**Affiliations:** Institute of Pharmaceutical Sciences, Pharmaceutical Chemistry, University of Graz, Austria; Institute of Biochemistry and Center for Molecular Biosciences, University of Innsbruck, Austria; Max-Planck Institute for Molecular Genetics (MPIMG), Otto-Warburg-Laboratory, Berlin, Germany; Tyrolean Cancer Research Institute (TKFI), Innsbruck, Austria; BioTechMed-Graz, Austria; Field of Excellence BioHealth - University of Graz, Austria

**Keywords:** protein interaction perturbation, deep scanning mutagenesis, MAVE, yeast two-hybrid

## Abstract

NF2 (Neurofibromine 2, merlin) is frequently inactivated in cancer, where its NF2 tumor suppressor functionality is tightly coupled to protein conformation. How NF2 conformation is regulated and how NF2 conformation influences tumor suppressor activity is a largely open question. Here we systematically characterized three NF2 conformation-dependent protein interactions utilizing deep mutational scanning interaction perturbation analyses. We identified two regions in NF2 with clustered mutations which affected conformation dependent protein interactions. NF2 variants in the F3 subdomain and the α3H helix region substantially modulated NF2 conformation and homomerization. Mutations in the F3 subdomain altered proliferation in three cell lines and matched patterns of disease mutations in neurofibromatosis. This study highlights the power of systematic mutational interaction perturbation analysis to identify missense variants impacting NF2 conformation and provides insight into NF2 tumor suppressor function.

## Introduction

NF2 (Neurofibromine 2, Merlin) is a member of the band 4.1 superfamily of proteins and is closely related to ERM (Ezrin-Radixin-Moesin) proteins, which functions as links between the cell membrane and actin filaments [1, 2]. While the other ERM protein family members do not have activities directly linked to cancer, NF2 tumor suppressor activity was initially characterized in flies and mice [3, 4], and later confirmed in mammalian cell models. NF2 links signals from the cell membrane to growth-related gene expression and acts in cell-cell contact inhibition [5–7], a function defined as one of the hallmarks of cancer. In contrast to the other ERM family members, NF2 lacks an F-actin binding domain. It also binds to phosphatidylinositol lipids [8]. NF2 is found at the cell membrane in contact with CD44, where it organizes cell junctions [9] and growth factor receptors [7, 10]. As an important regulator of cell growth, NF2 impacts proliferation associated pathways such as MAPK, AKT and Rac signaling [11, 12]. It also directly modulates transcription cofactor regulation, via AMOT/LATS or DCAF1, in the YAP/TAZ-hippo pathway [13–16]. More recently, NF2 was found to be an upstream factor of nucleic acid sensing, suppressing cGAS-STING-initiated antitumor immunity in cancer cell models [17].

Genetics also defines a prominent role of *NF2* loss of function in cancer. Genetic mutations or deletion of *NF2* cause neurofibromatosis type 2, an autosomal dominant disease predisposing to the formation of benign tumors. Biallelic *NF2* mutations cause tumor formation in the nervous system represented by vestibular schwannomas, meningiomas and ependymomas, frequently accompanied by hearing loss, dizziness, and neuropathies [18, 19]. *NF2* mutations are also commonly found in aggressive malignant mesothelioma, and frequently observed in other cancer types such as melanoma, breast, prostate, liver, and kidney cancers [20, 21]. *NF2* is a general tumor suppressor and cancer driver gene affected by mutations (tumor suppressor gene score 89% in pan cancer analysis [22]).

NF2 protein is expressed in various splice-isoforms with the canonical isoform 1 being the longest (**Figure 1a and S1a**). Isoform 2 has the same N-terminal part as isoform 1 and differs from isoform 1 only at the C-terminus through including exon 16 instead of exon 17. Ending with amino acid sequences from exon 16, the C-terminus of isoform 2 and 7 is five amino acids shorter than in isoform 1. In comparison to isoform 1 and 2, isoform 7 skips exons 2 and 3 and therefore lacks N-terminal amino acids 39-121 corresponding to most of the F1 and the initial amino acids of the F2 FERM subdomain.

**Figure 1:**
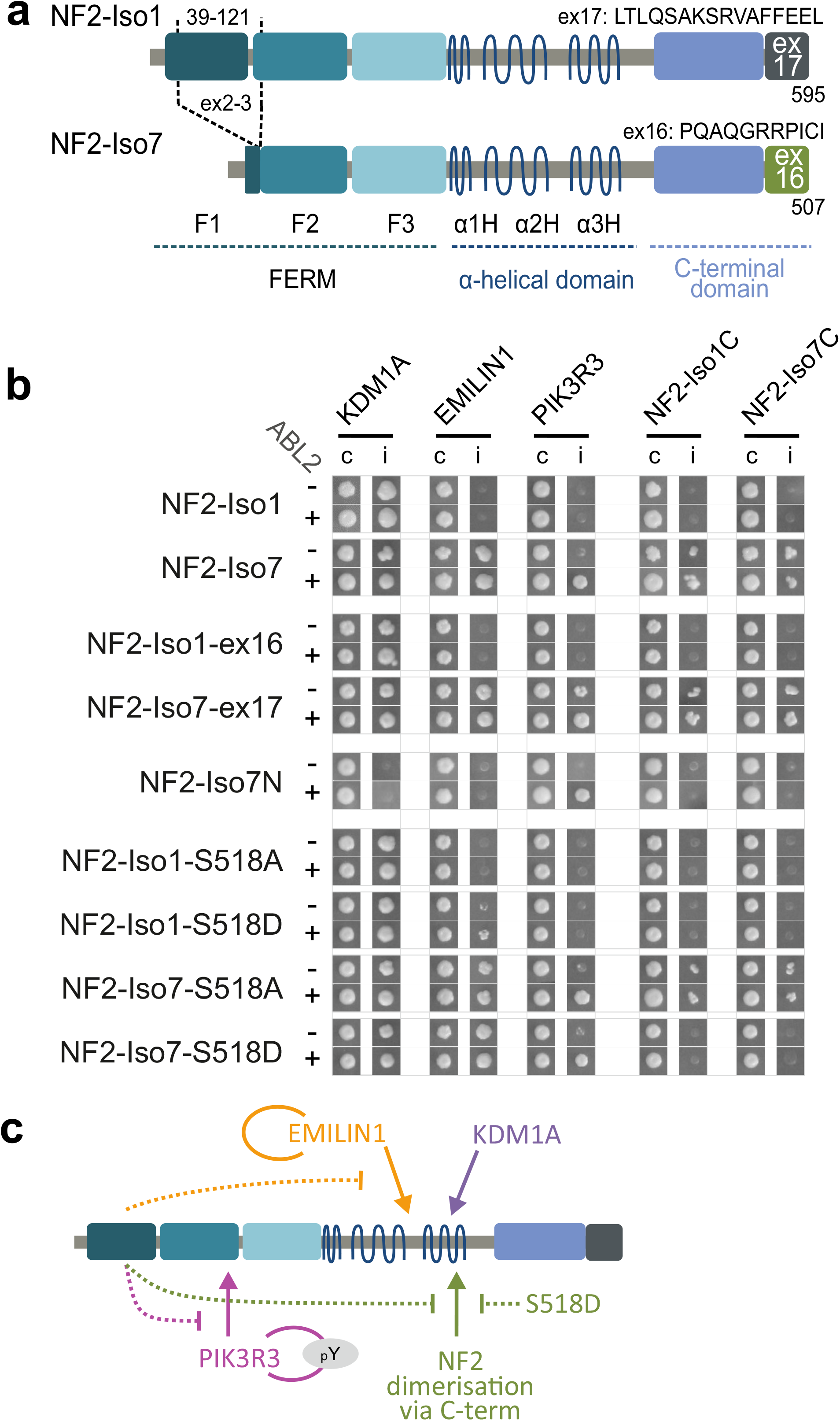
Conformation dependent PPIs of NF2. **a**. NF2 splice isoforms. The canonical isoform 1 is 595 amino acids long, the C-terminus is derived from sequences of exon 17 (595 AA [ex17], P35240-1). NF2 Isoform 7 is a shorter splice isoform (507 AA [Δex2/3, ex16], P35240-4). The C-terminus is alternatively spliced and includes exon 16 derived sequences which are 5 amino acid shorter (iso-1 580-590: LTLQSAKSRVA → PQAQGRRPICI, iso-1 590-595 missing), Isoform 7 lacks the majority of the F1 FERM subdomain and the beginning of the F2 subdomain encoded in exons 2 and 3, amino acids 39-121. Tumor suppressor activity has been demonstrated for isoform 1 but not for isoform 7. **b**. Y2H protein interaction results. NF2 bait constructs (row) were tested with five different prey constructs (columns) in the presence and absence of active tyrosine kinase ABL2 (+) and ABL2KD (K35M) (-) respectively. Prey: KDM1A (O60341), EMILIN1 (Q9Y6C2), PIK3R3 (Q92569), NF2-Iso1C (P35240-1: 308-595), NF2-Iso7C (P35240-4: 225-507). Growth of diploid yeast on non-selective agar (c) and on selective agar indicating protein interactions (i) is shown. **c**. Graphical summary of the interactions observed. Interactions with NF2 are shown with arrows, conformational interaction inhibition is indicated with dashed lines.

The isoform 1 has tumor suppressor activity while this activity was not demonstrated for all other NF2 isoforms including isoforms 2 and 7. Expression of isoform 2 was insufficient to impair growth in RT4-D6P2T cells [23, 24], however in HEI-193 cells stable expression of isoform 2 resulted in a reduction in relative BrdU-levels that was comparable to isoform 1 expression [25]. It was shown that deletion of exon 2 or exons 2-3 resulted in a dislocation of NF2 from the plasma membrane [26, 27]. A transgenic mouse model revealed promotion of Schwann cell proliferation by a NF2 mutant lacking amino acids 39-121, which was not observed for a C-terminal truncated NF2 isoform 1 protein version [28]. In *D. melanogaster*, deletion of a seven amino acid long stretch in F2 (AA 177-183 in HsNF2) was dominant negative for NF2 growth control [29] and a mutant construct with these seven amino acids replaced to alanine resulted in transformation and uncontrolled proliferation of cultured murine fibroblasts [30]. A splice isoform lacking exons 2-4 detected in hepatocellular carcinoma (HCC) cell lines was unable to suppress cell proliferation [31]. Based on the literature, we conclude that isoform 1 has tumor suppressor activity, while isoform 7 likely represents a version lacking tumor suppressor activity (**Figure 1a** and **S1**).

The importance of protein conformation for the tumor suppressor function of NF2 was recognized in early experiments [23]. Analogous to other ERM proteins, conformational rearrangements within NF2 are associated with functional changes, which are presumably triggered by post-translational modifications (PTMs), lipid- and protein-binding. NF2 can switch between an open and a closed conformation by self-association. The C-terminal residues of NF2 and a fully folded FERM domain are thought prerequisite for the formation of the head-to-tail interaction [32, 33], membrane localization [34] and tumor suppressor function [23]. However, alternative conformations have been proposed as well and the exact mechanism how NF2 conformation is regulated and how NF2 conformation influences tumor suppressor activity remains elusive [20].

For example, S518 is a critical phospho-site in NF2 [35, 36] and was suggested to influence signaling activity, localization, protein interaction and NF2 conformation. Most of the data suggest reduced tumor suppressive function caused by S518 phosphorylation. For example, in one study the phosphorylation mimicking S518D mutation in the full length and the C terminal construct prevented association with the N terminal construct, and the WT and S518A mutations allowed interaction with the N terminal part [37]. Conversely, in another study C-terminal NF2 constructs containing S518 phosphorylation mimicking mutations S518D and S518E increased the binding to N-terminal protein fragments while the S518A amino acid exchange abolished an intra-molecular interaction in co-IP experiments in HEI 193 cells [25]. Alternative models considering various degrees of conformational open- and closeness, are put forward to reconcile a series of apparently contradicting results [25, 38]. The interpretation of the data is complicated by experimental difficulties to monitor protein conformation and to clearly distinguish intra- and inter-NF2 interactions. NF2 homodimerizes at various degrees [32, 39, 40], and recently it was shown that gain of function mutations can even promote the formation of NF2 cellular condensates in conjunction with IRF3 [17]. Elucidating NF2 conformation and its effect on interaction partners and protein function remains a pivotal research task to better understand NF2 tumor suppressor activities.

Mutational scanning (DMS) has emerged as a powerful approach to systematically map amino acid residue-activity landscapes of proteins under defined readouts, yielding insights into protein function, structure, and evolution. At the same time these data assist computational variant effect prediction and clinical variant interpretation [41]. Physical interactions between proteins are critical to perhaps all biological processes and as such PPIs are basic cellular functional units that can be assayed universally for, in principle, all proteins [42, 43]. We developed a deep mutational scanning protein interaction perturbation screening technique based on reverse yeast two-hybrid (Y2H) analysis [44]. Here we used this method to scan a comprehensive set of single amino acid NF2 variants using four conformation dependent protein-protein interactions as readout. This allowed to investigate the mutational impact on NF2 protein conformational regulation and thus revealed amino acid residues critical for NF2 function.

## Results

### Conformation dependent NF2-protein interactions

We tested tumor suppressive NF2 isoform 1 and the shorter non-tumor suppressive isoform 7 (**Figure 1a**) for protein interactions and found three isoform-specific NF2 protein-protein interactions in a phospho-Y2H screen. Phospho-Y2H assays involve an active protein kinase delivered on a third plasmid in addition to bait and prey proteins, to also detect protein interactions that are modulated by phosphorylation of one of the interaction partners [45, 46]. The Y2H experiments revealed that Lysine-specific histone demethylase 1A, KDM1A, elastin microfibril interfacer 1 protein, EMILIN1, and phosphoinositide-3-kinase regulatory subunit 3, PIK3R3, interacted with NF2 (**Figure 1b**). The NF2-KDM1A [47–49] and NF2-EMILIN [48] interactions were listed in other systematic large scale PPI studies, however were not characterized any further. KDM1A equally interacted with NF2 isoform 1 and isoform 7, EMILIN1 interacted strongly with NF2 isoform 7, but not with isoform 1. Both PPIs were not affected by the presence of an active or a kinase dead version of non-receptor tyrosine kinase ABL proto-oncogene 2 (ABL2). In the case of PIK3R3, co-expression of active ABL2 was required for the interaction with NF2 isoform 7. PIK3R3 formed homodimers, and notably this PIK3R3 homodimerization was a pY-dependent interaction (**Figure S1b**). Therefore, the requirement of an active tyrosine kinase for the NF2-PIK3R3 interaction may be explained through a requirement for PIK3R3 dimerization. The NF2-PIK3R3 interaction was not observed with isoform 1 (**Figure 1b**).

We narrowed the NF2 binding site for the three interactions partners by using constructs that represented the N-terminal halves of isoform 1 (AA 1-K332, Iso1N) and 7 (AA 1-K249, Iso7N) and the C-terminal halves of isoform 1 (AA M308-595, Iso1C) and 7 (AA M225-507, Iso7C). EMILIN1 and KDM1A bound to the C-terminal half irrespective of the actual C-terminal ex17 or ex16 sequence (**Figure S1**). In contrast PIK3R3 interacted with the N-terminal part of NF2 isoform 7 (**Figure 1b** and **S1**). In summary, we found three isoform-specific protein interaction partner for NF2 through Y2H analyses - KDM1A, EMILIN1 and PIK3R3. When divided into two halves, the C-terminal part of NF2 protein interacted with KDM1A and EMILIN1, while PIK3R3 bound to a construct covering the isoform 7 N-terminal half of NF2. Because the longer isoform 1 contained all amino acids present in isoform 7, but does not interact with two of the three partners, we conclude that the protein interactions are sensitive to the NF2 conformation.

The very C-terminal residues of isoform 1 are thought to be necessary for the formation of a closed conformation [23, 32, 33]. Hence, we mutually exchanged the C-termini in isoform 1 and 7 and tested isoform 1 with an exon 16 derived - C-terminus (Iso1-ex16, identical to isoform 2) as well as isoform 7 with an exon 17 derived C-terminus (Iso7-ex17) for Y2H protein interactions (**Figure S1a**). Except for a weak increase in growth observed with Iso7-ex17 and PIK3R3 in the absence of active ABL2, NF2 protein interactions with mutually exchanged C-termini were the same as their wild-type counterparts (**Figure 1b**). This suggests that the isoform-dependent interaction specificity is not dictated by the very C-terminal part of NF2. Moreover, since EMILIN binds to the C-terminal half of NF2 (**Figure S1b**), the PPI pattern suggest that isoform specificity of the interaction is due to features in the FERM domain.

When we tested NF2 for homomeric interaction in the Y2H assay we found that full-length isoform 7 but not isoform 1 interacted with the C-terminal half of NF2 independently of whether the very C-terminal amino acids of the NF2 fragments or the full length NF2 partner resembled isoform 1 (ex17) or isoform 7 (ex16) (**Figure 1b**). The NF2 N-terminal constructs (Iso1N and Iso7N), when used as prey, did not show any interaction with full-length NF2. The NF2-Iso7N bait construct, while it interacted with PIK3R3, did not interact with either the Iso1C or Iso7C. In conclusion, this suggested, that the homo-dimeric interaction observed with the full length NF2 isoform 7 is mediated by the C-terminal part. Secondly, the isoform specificity had to be explained by the N-terminal part of NF2, because isoform-specific differences in the C-terminal amino acid sequence did not influence the NF2-NF2 interaction. Therefore, the NF2-Iso1 N-terminal FERM domain apparently causes the inhibition of a C-terminally mediated NF2-NF2 interaction, consistent with the hypothesis that isoform 1 adopts a protein conformation distinct to isoform 7.

As mentioned above different sets of experiments [25, 37], linked the phosphorylation of S518 to altered NF2 homomeric interaction and conformation using S518D phospho-mimicry and S518A phospho-null NF2 mutations. We tested the S518D and S518A mutant NF2 versions in Y2H assays for interactions with KDM1A, EMILIN1, PIK3R3 and NF2 itself (**Figure 1b**). The KDM1A interaction with NF2 was not substantially affected by changing S518 to either D or A. We observed a weak signal for the EMILIN - NF2-Iso1-S518D interaction and reduced growth for the PIK3R3 - NF2-Iso7-S518D pair in comparison to wild-type NF2. Interestingly, while the NF2 isoform 7 S518A mutant NF2 version, just as wild-type NF2-Iso7, did show homomeric NF2 interaction, S518D perturbed the NF2 interaction between mutated full-length isoform 7 and the NF2 C-terminal part (**Figure 1b**). Apparently phenocopying the isoform 1 FERM domain in NF2 in our assay, we concluded that the S518D phospho-mimicry mutant version negatively affects the PIK3R3 interaction and the NF2 isoform 7 homo-dimerization.

In summary (**Figure 1c**), our Y2H studies revealed that KDM1A interacted with NF2 isoforms 1 and 7, and EMILIN1 and PIK3R3 interacted with NF2 isoform 7 only. The PIK3R3 interaction was promoted through pY-dependent PIK3R3 homodimerization. We observed a homo-dimeric interaction between full-length isoform 7 and the C-terminal part of NF2 which was inhibited through either an isoform 1 FERM domain or a S518D phospho-mimicry mutation in full length NF2. It is important to emphasize that isoform 1 was active in the Y2H assay and contained all features of the shorter isoform 7. Since only the shorter isoform interacted with EMILIN and PIK3R3 and formed homomeric interactions, the isoform differences must be explained by indirect effects such as intra- or inter-molecular interactions of NF2. We hypothesized that the conformation of NF2 is critical for the isoform specific protein interactions patterns. Therefore, the binding status of KDM1A, EMILIN and PIK3R3 reflects different NF2 conformations. Single amino acid point mutations that are critical for NF2 conformation may therefore be reflected in alterations of protein interaction patterns.

### Deep mutational scanning interaction perturbation analysis of NF2

We developed reverse Y2H strains that can be used to select non interacting protein variants from complex genetic libraries [44]. The NF2-Y2H interactions gave robust growth repression on media lacking adenine, prerequisite for stringent reverse selection (**Supplemental Figure S2a**). We performed deep mutational protein interaction perturbation analysis with NF2 isoform 7 and its four wild-type interacting proteins: KDM1A, EMILIN1, PIK3R3 and NF2 C-terminal domain. A mutagenic library of NF2 isoform 7 containing all possible single amino acid exchanges to alanine (A::GCT), lysine (K::AAA), glutamic acid (E::GAA) and leucine (L::TTG) was generated using a multi-step PCR-based deep mutagenesis approach with on-chip-synthesized oligonucleotides [50]. The NF2 mutagenic pool was subcloned in the bait Y2H vector and used for transformation of the reverse-Y2H strains, mated in duplicates with the four prey strains in the presence of ABL2 and ABL2-KD (**Supplemental Figure S2b**). Interaction perturbing NF2 variants for a given partner were enriched through growth on selective agar lacking adenine and disruptive mutants were identified through next-generation-sequence analysis.

Statistical data analyses of the sequence reads from 6 controls (no interaction selection) and 36 mutant library interaction samples (each biological replicate was sequenced three times) resulted normalized interaction perturbation profiles of the NF2 isoform 7 with its four partners respectively (**Supplemental Figure S2c and Supplemental Table S1**). The mutant NF2 libraries did not show a mutation selection bias during the cloning and strain preparation procedures (**Figure 2a**) and covered the whole NF2 sequence uniformly (**Figure 2b**). Efficient enrichment of programmed AKEL mutants through interaction selection with the rY2H system was observed, when comparing the input library to sequences obtained from the mutant library tested against a wild-type interaction partner (**Figure 2c**). For all NF2 interaction pairs, we observe a clear deviation from the codon and position specific linear models for the AKEL mutations but very little deviation for all other sequenced mutations (**Figure 2d**).

**Figure 2:**
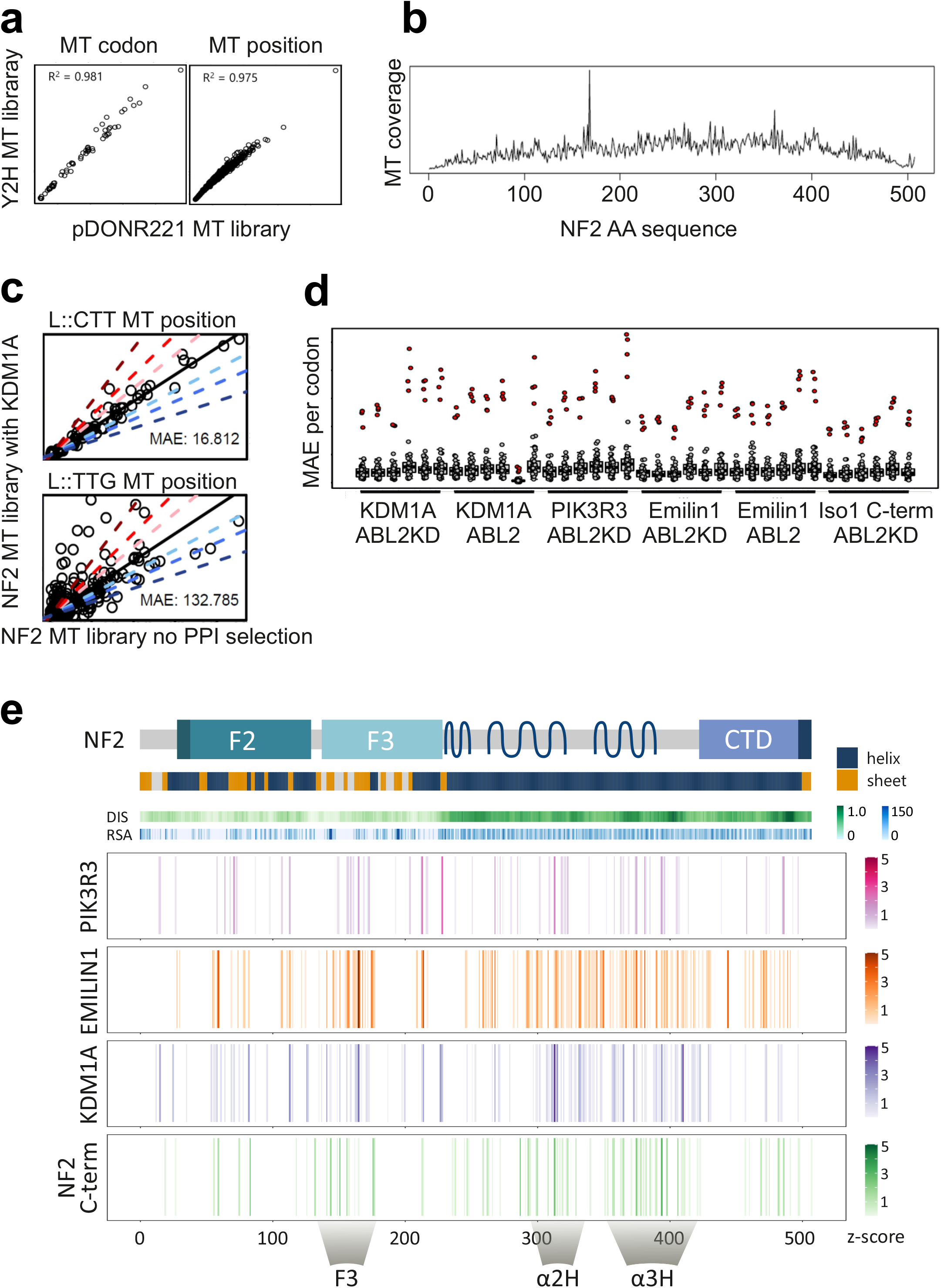
Deep mutational NF2 interaction perturbation. **a**. Number of mutants reads from the mutant NF2-Iso7 library in pDONR221 (library cloning vector) and pBTM116-D9 (Y2H vector). The axes represent the number each mutation was sequenced for a codon (left) or for a NF2 position (right). **b**. Mutational sequence coverage (number of mutant reads) of the NF2 mutant library across the NF2 amino acid sequence. **c**. Example of the recall statistics of the rY2H selection with NF2-KDM1A Enrichment of a non-programmed mutant codon (L::CTT) and a programmed mutant codon (L::TTG, right) after rY2H selection of mutant library NF2 isoform 7 through interaction with wildtype KDM1A across all positions within NF2. Enrichment of reads in comparison to the NF2 MT library for a set of programmed (L::TTG) mutations is observed as deviation from a linear model (mean absolute error MAE = 132). **d**. Overall results for all PPI samples. Deviation from codon specific linear models (MAE: PPI vs NF2 library) of the programmed amino acid mutations (red dots) is much larger than of any other mutations (grey dots). MAE values of all codons of the for all 36 protein interactions sequenced are shown. **e**. rY2H interaction perturbation profiles. Schematic of NF2-Iso7 protein, domain structure with structure (helix/sheet), disorder prediction (DIS) and solvent accessibility predictions (RSA). Aligned, combined enrichment profiles of the four interactions are shown. Sequence position interfering with the NF2-Iso7 interaction are color coded (Z-score). The PIK3R3 profile represents a combination of two biological replicas assayed in the presence of ABL2, the NF2 C-term profile combines two experiments with ABL2KD. KDM1A and EMILIN1 profiles contain a combination of four biological replicas (each two with ABL2 and two with ABL2KD). Mutational cluster regions are highlighted as F3, α2H and α3H.

Enrichment values were normalized across the interactions and expressed as Z-score. Even though we located the respective interaction sites of the interaction partners differently to the N- or C-terminal halves of NF2, disrupting mutants were found spread over large parts of NF2 for all four interaction partners (**Figure 2e**). This result likely reflected both, mutations that disrupt folding as well as mutations linked to conformational constraints of NF2. However, three hotspot regions emerged from the interaction perturbation patterns of the interactions. Amino acid substitutions in the N-terminal half were selected perturbing the interactions in an area in the F3 part of the FERM domain. In the C-terminal half of NF2, we found a region with clustered mutations in the α2H helix (around AA 310 in isoform 7), and a second region in the α3H helix (proximal to AA 350-400 in isoform 7). The interaction perturbation profiles of the four tested interactions (**Figure 2e**) showed substantial overlap in three critical regions located in the N-terminal and in the C-terminal half of NF2.

### Assessing single site mutations in NF2-interactions

We next validated the deep mutational scanning results in order to identify mutations that selectively alter protein interactions without affecting overall protein folding and stability. 60 amino acid positions were selected and mutations were individually introduced in both NF2 isoform 1 and 7. Constructs were confirmed through sequencing and used for individually testing each interaction with the WT interaction partner proteins in a pair-wise Y2H colony spot matrix assay. 16 of the single-site mutations caused a selective change of interaction patterns compared to interactions observed in their WT isoform counterparts (**Figure 3a**). We observed distinct patterns of loss and gain of interactions with the two NF2 isoforms and its four interaction partners (**Figure 3b**). 5 single amino acid point mutations altered the NF2 interaction with KDM1A, 4 with EMILIN, 15 with PIK3R3 and three substitutions modulated the NF2-NF2 interaction.

**Figure 3:**
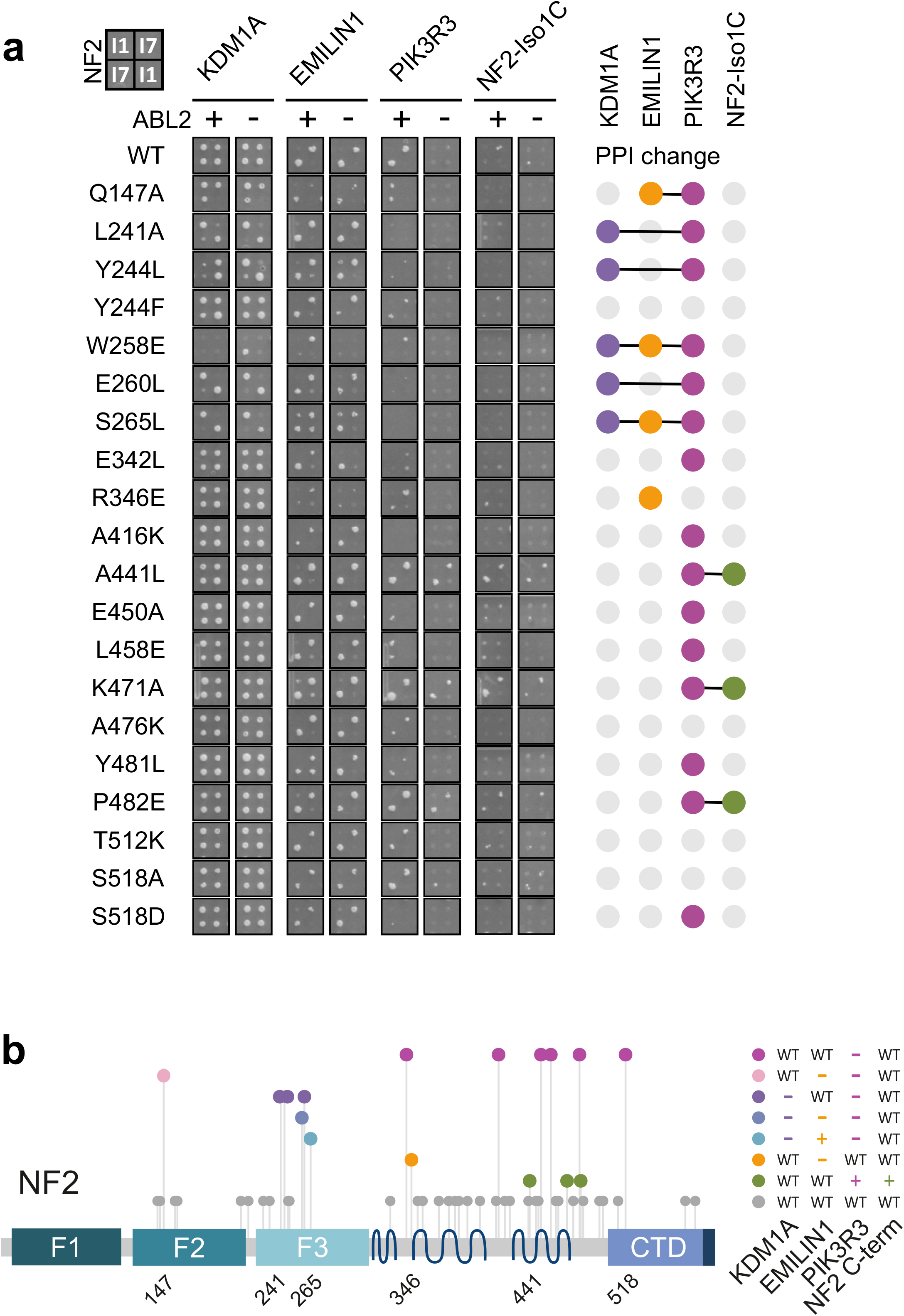
Distinct patterns of loss and gain of interactions with NF2 variants. **a**. Y2H protein interactions of a selected subset of NF2 single-site mutants. Selective agar, where yeast colonies indicate protein interactions, NF2 bait constructs in rows and interacting prey in columns. Each individual mutation (isoform 1 numbering) was tested as isoform 1 (384 plate format: upper left and lower right spot) and isoform 7 (384 plate format: upper right and lower left spot) in duplicate in the presence of an active tyrosine kinase ABL2 (+) or an inactive version ABL2KD (-). Right: upset plot indicating alterations of the interaction patterns in comparison to wild-type NF2 through colored dots. **b**. Mutant classes defined in the Y2H spot assay. The Y2H validation assay with single-site mutations performed in NF2 isoforms 1 and 7 resulted in specific interaction losses and gains when compared to the two WT isoforms, respectively. Projected to the protein primary structure of NF2, mutants were grouped into eight classes, including wild-type (grey) and four mutations with individual PPI patterns.

The largest class of mutants behaved like WT NF2 in the Y2H assay which can be explained by the different sensitivity in the deep scanning screen compared to pair wise spot testing. Four mutations, Q147E, S265L, R346E and W258E (isoform 1 numbering used throughout), showed unique patterns of interaction perturbation (**Figure 3b**) where W258E impacted all three PPIs. Q147E reduced PPIs with EMILIN and PIK3R3, R346 resulted in a selective loss of EMILIN interaction and the Iso1-S265L variant increased the EMILIN interaction. Interestingly, L241A, Y244L, E260L and S265L reduced the KDM1A interaction selectively with the isoform 7, while the substitutions did not affect the PPIs with isoform 1. PIK3R3 did not interact with these variants any more, but the interaction of the four isoform 7 variants with EMILIN was not altered.

A large group of variants represented specific PIK3R3 interaction disrupting mutations distributed across the whole protein sequence (**Figure 3a**, magenta). As defined in the rY2H-seq screen, the majority (A416K, E450A, L458E, Y481L) localized in the C-terminal clusters α2H and α3H. In the colony assay, the interaction of NF2 with PIK3R3 and with NF2-Iso1C appeared generally weaker. This allowed the observation that three α3H mutations, A441L, K471A and P482E strengthened the PIK3R3 interaction rendering it independent of ABL2 dependent PIK3R3 dimerization. The same NF2 variants also resulted in a strengthened homomeric interaction of NF2 isoform 7 with the NF2 isoform 1 C-terminus (**Figure 3a**, green). Comparable to S518D, which negatively affected both the interaction with PIK3R3 and the NF2-NF2 interactions, the three α3H mutations also showed a coupled phenotype promoting both the PIK3R3 interaction and the NF2-NF2 homomerisation.

Notably, in the colony assay, all five KDM1A interaction perturbing mutations were clustered in the F3-FERM subdomain. This is intriguing given the fact that our Y2H experiments showed that the C-terminal half of NF2 was sufficient to interact with KDM1A. The result can be explained by an indirect, conformation driven perturbation of the interaction caused by the F3-FERM domain mutants. In summary, we identified at least two novel regions important for NF2 regulation of its protein interactions and conformation. The N-terminal F3 region in the FERM domain is a key determinant to isoform interaction specificity and the α3H in the C-terminal half of the protein is involved in NF2-NF2 interactions.

### Structural impact of FERM domain mutations

Mapping the C-terminal mutations in α3H which enhanced the NF2-NF2 interaction (A441L, K471A, P482E) to a structural prediction model of full length NF2 isoform 1 (AF-P35240-F1) revealed a localization of the residues in extended alpha helical rod-like structures of the protein (**Supplemental Figure S3**), providing limited information for interpretation. However, we mapped the mutations in the N-terminal half of NF2 to a 3D atomic structure of the FERM domain solved in a closed NF2 protein state (PDB: 4zrj, [51]). FERM domain structures contain three subdomains (F1, F2, F3) forming a cloverleaf structure (**Figure 4a**). The F1 subdomain resembles ubiquitin, F2 has similarities to the acyl-CoA binding protein and the F3 subdomain has structural similarities to phospho-tyrosine binding, pleckstrin homology, and Ena/Vasp Homology 1 (EVH1) signaling domains. We note that this closed 3D structure also includes a peptide from the NF2 C-terminal part with two A585W/S518D stabilizing mutations (chain B, 506-595) where the tryptophan 585 inserts into the F3-FERM domain (**Figure 4a**). In silico mutagenesis (BIOVIA Discovery Studio 2020) allowed to model potential effects of mutations on the conformation of NF2. For example, Q147A, which had a uniquely altered interaction pattern, is located in the F2 part of the FERM domain. The wild-type Q147 formed four hydrogen bonds (<3.4 Å), two of them connected to E152 and R198 in other helixes within the F2 domain (**Figure 4b**). Upon replacing Q with A in the structure, the hydrogen bonds to neighboring alpha helices were disrupted and weak hydrophobic interactions formed.

**Figure 4:**
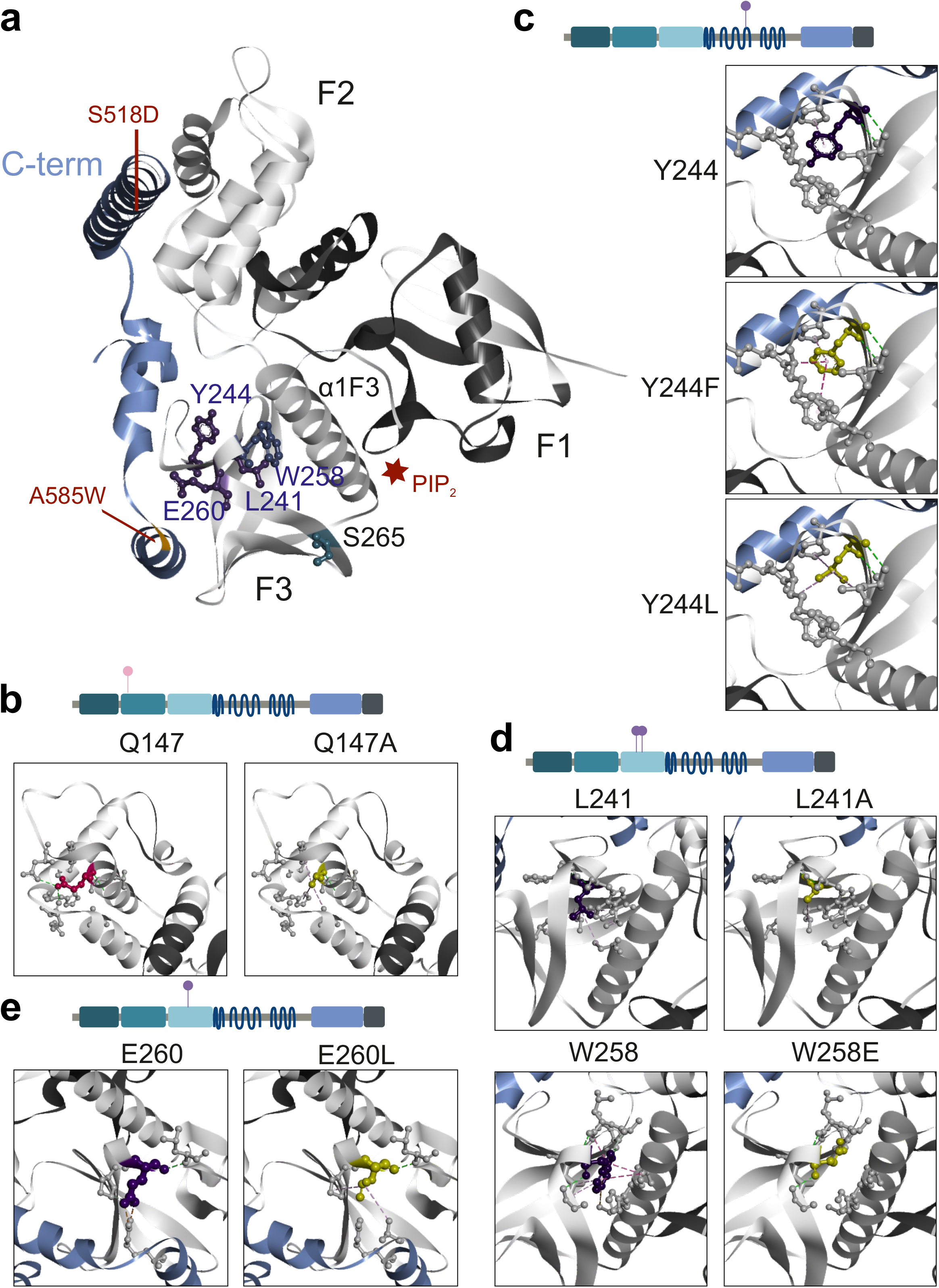
Structural analyses of NF2 variants. **a**. FERM domain mutations projected on the NF2 protein structure. 3D ribbon “cloverleaf” structure of the FERM domain in grey, C-terminal NF2 peptide in blue colors (PDB: 4zrj). Mutations S518D and A585W in the C-terminal NF2 peptide that stabilized the structure are indicated, as well as the PIP2 binding groove between F3 and F1 subdomain (red star). Amino acid positions perturbing the NF2 interaction pattern (purple, ball and stick) cluster in the F3-FERM subdomain. **b-d**. 3D structures of the FERM domain as ribbon models in which indicated residues are colored and displayed as ball and stick atomic model with the respective interactions indicated as dashed lines. Wild-type residue (left panel) in comparison to mutant residue (right panel). **b**: Q147; **c**: Y244, **d**: L241 and W258, **e**: E260. The models were created with BIOVIA Discovery Studio 2020 (version 20.1.0.19295).

We propose that the mutations at positions Y244, L241 and W258, which spatially clustered in the central part of the F3 lobe of the FERM (**Figure 4a**), may perturb positioning of the large alpha helix (α1F3) in the F3-F1 interface thereby affecting overall NF2 conformation. Residue Y244 was located very central in the F3 subdomain and had a hydrogen bond with amino acid L233 (∼3 Å) and one hydrophobic interaction with P252 (4.2 Å) (**Figure 4c**). A Y244F mutation did affect these bonds, yet added weak hydrophobic interactions to Y221 (5.2 Å) and R249/L250 (4.7 Å) were observed. Modeling a Y244L mutation added weak hydrophobic interactions with R249 (5.5 Å) and L233 (5.0 Å).

The intramolecular interactions of L241 involved one hydrogen bond to F256 and four additional hydrophobic interactions with amino acids L234, V236, I273, C300 (≥ 4.7 Å). The model with a L241A exchange lacks all hydrophobic interactions except L234(5.4 Å, **Figure 4d**). The loss of contacts affected interactions with the neighboring β4F3-sheet and the α1F3-helix, the helix connecting the F3 subdomain with the central helical domain. We made a similar observation when modeling the W258E substitution (**Figure 4d**). In the 3D structure W258 forms hydrogen bonds to residues D237, A238, L239 and I261 (∼3.0 Å), hydrophobic interactions with N303/H304, L239/G240, L241, I261 and a Pi-Sulfur interaction with C300. A glutamic acid at position 258 lost all contacts that connected the W258-β-sheet with the α1F3-helix. The structural analysis suggested that mutations at both positions, L241 and W258, perturb the F3 domain interaction with the α1F3-helix. The α1F3-helix was shown to undergo a large structural rearrangement upon PIP_2_ and LATS binding [52, 53] (**Figure 4a**).The structural modeling of mutations, although not dynamic, can provide hypotheses which address these NF2 sites within the F3 domain as key residues influencing the conformational state of NF2.

E260 spatially clusters with the other interaction perturbation F3-FERM domain residues (**Figure 4a**), however it does not point towards α1F3-helix, but in the opposite direction. E260 is actually at the contact interface of the F3-FERM subdomain and the C-terminal end of NF2 (**Figure 4e**). E260 forms a strong salt bridge to R588 localized in the C-terminus of NF2 (2.7 Å). Upon modeling a mutation to leucine at the position, this salt bridge was perturbed and replaced with a 4.9 Å hydrophobic contact. Hydrogen bonds between E260 and P257 and L276 within the F3 domain were hardly affected. For modeling the E260L mutation, the Alphafold protein structure prediction (AF-P35240-F1) had to be utilized as E260 is in close proximity of the A585W stabilizing mutation in the 3D Xray structure (4zrj) (**Figure 4a**). The tryptophan at 585 sterically interferes with modelling the E260L exchange, hence this further suggests that E260 is at an important place for contacts across NF2 to the C-terminal part.

In summary, the structural analysis of NF2 variants with altered interaction patterns showed that these substitutions affected contacts to regions distant in the primary sequence, either within the FERM domain or to the C-terminal part of NF2. This observation supports the hypothesis that the sites are critical for the structural dynamics of NF2 and defines the F3-FERM domain, in particular through positions, L241, W258, and E260, as critical structure for the overall NF2 protein conformation.

### Effect of NF2 variants on cell proliferation

We examined NF2 variants for their cellular effects on proliferation in mammalian cell culture. The FERM domain is critical for subcellular NF2 protein localization [34]. For example, isoform 7 delocalized from the plasma membrane to cytoplasmic structures [26]. A crystal structure of NF2 bound to PIP_2_ highlighted the grove between the FERM F1 and F3 subdomains as phospho-lipid binding site in NF2. Lipid binding deficiency and altered protein interactions may be coupled [52]. Therefore, we first examined subcellular localization of the NF2 FERM domain mutant variants in fluorescence microscopy (**Figure 5a**). Using transient transfection of YFP fused-NF2 mutant constructs in HEK293T cells, we did not observe pronounced differences with respect expression levels and subcellular localization compared to the wild-type NF2. NF2 was mainly found at the plasma membrane, especially at cell-to-cell contact sites, and also in the cytosol. Partial membrane localization was observed for all variants. Iso1-W258E, which was expressed to similar levels as wild-type and the other mutant NF2 proteins, however was more concentrated in the perinuclear region.

**Figure 5:**
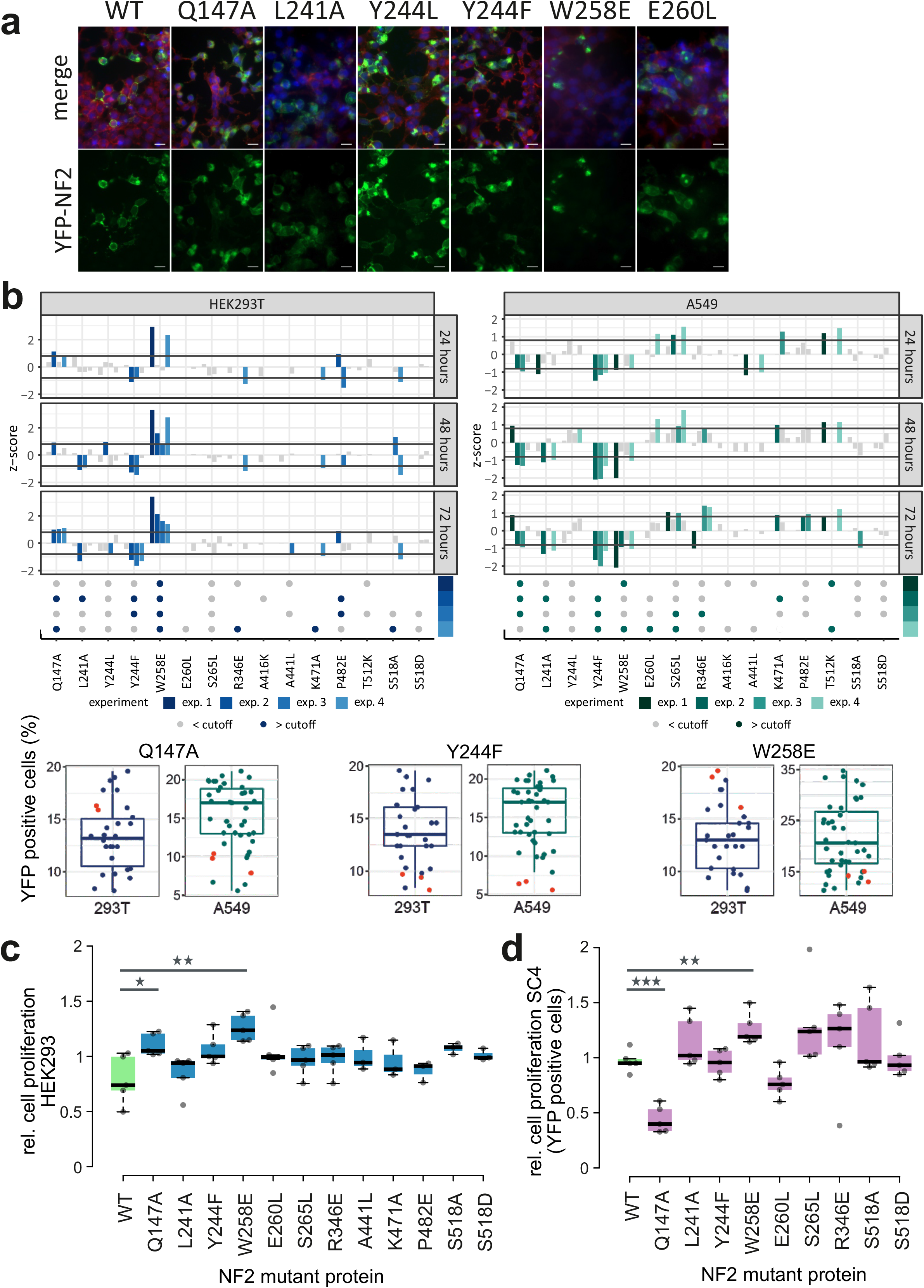
Impact of NF2 variants on cell proliferation. **a**. Fluorescence microscopy images of wild-type and NF2 variants. HEK293T cells were transiently transfected with N-terminal YFP-tagged NF2 (green) and cellular localization of NF2 proteins 24h after transfection was investigated by fluorescence microscopy. Cells were fixed, permeabilized and nuclei stained with DAPI (blue). Actin filaments were stained with Rhodamine-Phalloidin (red). Top, merge of all three stains, bottom, YFP signal from NF2 expression. Scale bar = 20μM. **b**. FACS based proliferation analyses of HEK293T and A549 cells expressing YFP-tagged NF2 proteins. Cells were transiently transfected with a plasmids expressing YFP-tagged NF2 WT and mutant versions and the relative fraction of YFP positive cells was determined over a time period of three days. Individual experiments were performed as triplicates, each NF2 mutant versions was tested in up to four experiments. Upper panel: Z-scores were calculated and variants below or above the cutoffs of -0.8 or 0.8 were colored according in each experiment. Dots: constructs which exceeded the cutoff on two individual timepoints in one experiment were colored in blue, variants below the cutoff are colored in grey. Lower panel: Examples of individual experiments where the fraction of YFP-positive cells at 48h is shown. Data points of the Q147A, Y244F and W258E NF2 variants are shown as red dots respectively, blue dots show the distribution of all other NF2 protein variants in the experiment. **c**. Live cell imaging of HEK293 transiently transfected with N-terminal YFP-tagged NF2 variants. Relative cell proliferation is calculated as area of cell confluency over select time intervals between 16h and 64h after transfection and normalized to the median cell proliferation of all NF2 constructs. Box plots represent the average of triplicate measurements of five experiments. Statistically significant differences (t-test, * = p < 0.05; ** = p < 0.01) to wild-type NF2 are indicated. **d**. Live cell imaging of SC4 NF2^-/-^ cells transiently transfected with N-terminal YFP-tagged NF2 variants. Relative cell proliferation is calculated as area of cell confluency of YFP positive cells over select time intervals between 12h and 40h after transfection and normalized to the median cell proliferation of all NF2 constructs. Box plots represent the average of triplicate measurements of five experiments. Statistically significant differences (t-test, ** = p < 0.01; *** = p < 0.001) to wild-type NF2 are indicated.

We next assayed the effects of the single amino acid substitutions on cell proliferation, a key feature of the tumor suppressor function of NF2 [25, 52–54]. Fluorescence-activated cell sorting (FACS) analysis was performed to investigate the number of transfected mutant NF2 YFP-positive cells every 24 h over a 72 h period in four experiments (triplicates each). The percentage of YFP positive cells was recorded and z-scores were calculated for each of the 15 mutant constructs in each experiment and timepoint. Because apparently contradicting results on NF2 functional effects may be attributed to the use of different cell lines [20], we tested both, HEK293 (epithelial human embryonic kidney) and A549 (epithelial lung carcinoma) cell lines. Both cell lines allowed for a relative high rate for transient transfection and are frequently used for proliferation assays.

HEK293 and A549 showed differences in their overall response to NF2 transfection. While the fraction of HEK293 wild-type NF2 transfected cells (YFP positive) decreased over time, this was not observed in the A549 cell line. However, we investigated every cell line separately and assessed the effect of mutant NF2 transfection versus the wild-type protein. Across the 15 NF2 variants tested (S518A and S518D included) three FERM domain substitutions had the strongest and best reproducible effect on the fraction of YFP positive cells (**Figure 5b**). In HEK293, Y244F showed a relatively lower YFP-positive cell number, Q147A and W258E had a relative higher YFP-positive cell fraction compared to the group of all NF2 variants. Although the A549 cell line had a higher inter-experiment variation in the FACS quantifications, Q147A, Y244F, W258E and S265L significantly affected proliferation. Q147A, W258E showed opposite effects in the two cell lines, which may be attributed to the different genetic background and cell line specific proliferation requirements. Consistently across the two cell lines, our analysis showed that mutations altering NF2 conformation, specifically Q147A, Y244F, W258E in the F3 region, affected cell proliferation.

To confirm the FACS analyses, we monitored cell proliferation of YFP-tagged NF2 variants through live cell imaging using a Incucyte phase and fluorescence imaging analysis system. We transfected HEK293 cells with the wild-type and mutant constructs, tracked cell proliferation by quantifying cell confluency over time. In five experiments, relative cell proliferation was normalized to the average cell proliferation of all NF2 transfected cells. Confirming the results from FACS analysis with HEK293, wild-type NF2 reduced cell proliferation and Q147A and W285E significantly perturbed this effect (**Figure 5c**). Finally, we performed an analogous analysis in SC4 cells, an immortalized mouse schwannoma NF2^-/-^ cell line, that has been used to study NF2 proliferation effects [11, 12]. Because transfection efficiency of SC4 cells was about 10-20% only, we quantified YFP-positive cell area through live cell imaging of the NF2 variants in five experiments. Transfection of SC4 cells with NF2 Q147A and W258E significantly reduced and elevated cell proliferation in comparison to wild-type NF2, respectively (**Figure 5d**). The NF2 Q147E effect is in line with the results observed in A549 cells and W258E with HEK293 respectively. These experiments in mammalian cells demonstrate that the amino acid substitutions in the FERM domain alter cell growth in an NF2-dependent manner.

## Discussion

In analogy to ERM proteins, NF2 function and tumor suppressor activity is thought to be controlled by conformation switches between an open or a closed state. In the case of ERM proteins, the functionally activating, conformational change is mostly driven by phosphorylation of a threonine T567 residue in the C-terminal domain, a mechanism that as such does not exist for NF2 [1, 55]. Whether or not post-translational modifications of NF2, for example at S518, cause any substantial conformational changes appears to be context specific [25, 38, 56, 57]. While NF2 conformation determines its function, it remained unclear which conformation is active and what determines the NF2 conformation upon activation [20].

Structural data obtained with a set of protein interaction partners, highlight conformationally different NF2 states induced by protein- and lipid binding [52, 53, 57]. For example, the NF2 interaction partner AMOT increased binding of NF2 to LATS1 [51]. Since the NF2 binding sites for LATS1 and AMOT were reported in the FERM domain (F2 subdomain) and the central helical domain (AA 401-550), respectively, it was concluded that AMOT binding induced a conformational opening of the FERM domain LATS1 binding site [51]. *In vitro*, LATS1 binding increased by 10-fold when NF2 was preincubated with PIP_2_, and lipid binding (in the F1-F3-cleft, **Figure 4a**) was suggested to release the C terminal-FERM domain intra/inter molecular NF2 contacts exposing the LATS1 binding site [52]. On the other hand, DCAF1 binding to the F3 domain had no major influence on NF2 conformation [58–60].

Homomeric interactions with various NF2 constructs were observed [32, 39]. Y2H experiments and co-immunoprecipitation assays showed full length isoform 1 dimerization with full length isoforms 1 and 2 [32, 39], and interaction of the C-terminus of isoform 1 and of isoform 2 [39]. N-terminal NF2 (AA 1-313) was used to precipitate full length NF2 isoform 1 *in vitro*, while the N-terminal part was not able to interact with NF2 isoform 2 [61]. N-terminal constructs interacted with full length NF2 and C-terminal constructs inY2H and *in vitro* binding assays [23, 25, 32]. Therefore, current models of NF2 dimerization allow both N- to C-terminal antiparallel interaction and C- to C-terminal interaction. Similar to the PPI patterns with PIK3R3, our experiments demonstrated C- to C-terminal NF2-Iso7 interactions that were indirectly regulated by the FERM domain in the N-terminal part of NF2. We cannot rule out other NF2 homomeric interactions, for example NF2 isoform 1 homodimers, which may have occurred as false negatives in our Y2H setup.

In total, 141 protein interaction partners of NF2, including KDM1A [47–49] and EMILIN [48], were previously annotated in systematic large-scale human PPI studies, most of which are not functionally characterized [62]. While we did not address the potential biological function of the NF2 interactions with KDM1A, EMILIN, or PIK3R3 in this study, we thoroughly characterized differential isoform specific, conformation-dependent interaction patterns with these three NF2 interaction partners. The NF2 interaction partners bound to different parts of the protein, however EMLIN, PIK3R3 and NF2-Cterm binding was negatively regulated by FERM domain structures distant in the primary sequence (**Figure 1c**). Our objective was to utilize the interactions to probe NF2 conformation in deep mutational interaction perturbation scanning experiments.

Deep mutational scanning is a powerful approach for determining the sequence-function relationships with the goal to better predict the functional consequences of genetic variation [42, 63], identify sites that regulate protein interaction [44, 64, 65] or probing protein conformation and structure [66]. Variants of other key tumor suppressor proteins, such as PTEN and BRCA1, were successfully characterized using mutational scanning approaches coupled to functional assays [67, 68].

In this study we have assessed the effect of > 2000 single amino acid substitutions in tumor suppressor NF2 on the interactions with four protein partners that bind NF2 in a conformation dependent manner. In our interaction perturbation approach, we mutated all NF2 position to either a glutamic acid (E) or a lysine (K) introducing a negative and positive charge, respectively. We also replaced the wild-type amino acids with alanine (A) as well as leucine (L) introducing a small and a large residue, respectively. Other exchanges such as to proline, glycine, tyrosine or tryptophan were found to frequently affect stability and expression level of the protein variants [65]. On one hand, the choice of using four different amino acids instead of all possible in our experiment may limit our inferences with respect to disease variants. On the other hand, we introduce more subtle perturbations to specifically probe interactions and protein conformation [44]. Loss of interaction can be caused by effects on binding affinity or on protein abundance (or both) [65]. We did not quantify these effects, but robustly controlled for protein folding effects in our experiments through the use of four distinct binding partners. Interaction selectivity of the perturbation effects (**Figure 3**) demonstrated that the studied mutations do not substantially affect protein abundance.

For all four interaction partners, we observed relatively wide spread mutation perturbation across the primary sequence of NF2 (**Figure 2**). The patterns are more similar than expected, and did not directly reflect larger primary binding sites of the partner proteins. Rather, both N- and C-terminal NF2 parts appeared as critical determinants for the NF2 interactions in line with your hypothesis that the PPIs reflect NF2 conformation. We observed multiple signals in the N-terminal part of NF2 when probed with EMILIN and KDM1A for which the C-terminal part was sufficient for binding. Conversely, multiple mutations in the C-terminal part affected PIK3R3 binding, for which we located the primary binding site in the N-terminal part. In our validation experiments, where we tested selected amino acids substitutions individually (**Figure 3**), 15 NF2 variants showed altered PPI pattern. Critical mutations clustered in two regions (**Supplemental Figure S3**). The F3-FERM subdomain harbored all five mutations modulating the KDM1A and, with the addition of R346E, the EMILIN interactions in the colony assay. In addition to the F3-FERM domain regions which appeared key to NF2 conformation, other clusters of substitutions in the α2H and α3H affected the PIK3R3 interaction. We showed in our pY-Y2H studies that PIK3R3 interacted as dimer with NF2. Three mutations in the αH3 helix (A441L, K471A, P482E) relieved the requirement for PIK3R3 dimer formation for NF2 interaction and also promoted NF2-NF2 C-terminal interaction. Therefore, we identified novel single site determinants, other than S518, important for NF2-NF2 interactions.

Studying the FERM domain mutations on the 3D domain structure (**Figure 4**) illustrates that positions Q147 Y244, L241, W258 and E260 form contacts bridging different secondary elements of the FERM domain most notably to helix α1F3. Modeling the respective mutations disrupted several of those contacts impacting the dynamics of the FERM domain and thus potentially allowing for large scale conformational changes. Indeed, the Q147A, Y244F and W258E NF2 variants most strongly affected cell proliferation in FACS and live cell image analyses (**Figure 5**). This result was consistent across three epithelial cell lines, HEK293, A549 and SC4, however the effects appeared context dependent. Different effects may – for example – be due to different NF2 endogenous levels, other endogenous NF2 interaction partners, the interplay with other tumor suppressor protein activities or distinct dependencies on growth pathways in the three cell lines. NF2 Q147A showed increased repression of cell proliferation in comparison to wild-type in A549 and SC4 cells, while W258E showed a decreased suppression of cell proliferation in HEK293 and SC4 cells. The latter finding is in agreement with the observation that W258E largely lost cell membrane localization and that delocalization from the membrane is a condition associated with loss of tumor suppressive function of NF2 [69, 70].

In addition to functional differences associated with NF2 versions lacking exons 2-3 [28, 31], several patient-derived type 2 neurofibromatosis NF2 missense mutations within the N-terminal FERM domain were found which impaired NF2 interactions with DCAF1 [14]. Missense L46R, L64P, or L141P mutations in the FERM domain were shown to convert NF2 into a loss of function phenotype, suppressing innate DNA sensing and STING-initiated antitumor immunity [17]. There were 430 NF2 missense mutations deposited in the archive of human genetic variants and interpretations of their significance to disease, ClinVar (Dec 19, 2021) [71]. 406 were annotated as variants of uncertain significance (VUS), 11 as conflicting and only 13 as either likely benign, likely pathogenic or pathogenic in type 2 neurofibromatosis. VUS included Q147P, Y244C, W258G, S265L, R346S/K, A416V, A441P, L458R, Y481C and P482R/L, amino acid residue positions for which we provide multiple evidence that substitutions are likely to impact NF2 conformation and function (**Supplemental Figure S3**). Therefore, our mutagenesis data aids NF2 variant interpretation. Additionally, a substantial set of NF2 variants was subjected to validation and functional testing providing novel mechanistic insight. Our data reveal two functional important regions for NF2 conformational dynamics. The α3H helix in the C-terminus mediates NF2 homomeric interactions which are critical for NF2 activity. The FERM domain, specifically the F3, appears as key trigger for conformational regulation of NF2 suppressor function. Our variants provide useful genetic tools for further mechanistic studies of context dependent NF2 function.

## Material and Methods

### Mutagenic library preparation

The mutagenic library of NF2 was generated using a multi-step PCR-based deep mutagenesis approach with on-chip synthesized oligonucleotides from Custom Array, Inc.. A total of 2145 NF2 primer sequences encoding NF2 single amino acid substitutions were synthesised. Each single amino acid was exchanged to alanine (A::GCT), lysine (K::AAA), glutamic acid (E::GAA) and leucine (L::TTG). The mutagenesis protocol was essentially carried out as in Kitzman et al, [50], with adaptations described in Woodsmith et al. [44].

### Interaction perturbation reverse Y2H screen

Reverse Y2H strains that can be used to select non-interacting protein variants from complex genetic libraries [44] were used for interaction mating [**RPrey_S3:** MATα, *his3-Δ200, trp1-901, ade2, leu2-3, 112, gal4, gal80, can1, cyh2, met2, ura3::KanMX::(lexAop)*_*8*_*-GAL1TATA-lacZ, LYS2::(lexAop)*_*4*_*-HIS3TATA-HIS3, met2::((LexAop8::TetR)*_*2*_*), ho::(LexA8::TetR)*_*2*_*-ura3;* **RBait_S3:** MATa, *his3-Δ200, trp1-901, ade2, leu2-3, 112, gal4, gal80, can1, cyh2, met2, ura3::(lexAop)8-GAL1TATA-lacZ, LYS2::(lexAop)*_*4*_*-HIS3TATA-HIS3, met2::(TetO5A::ADE2), ho::(LexA8::TetR)*_*2*_*-ura3]*. The **RBait_S3** MATa strain was transformed with the mutant libraries according to a lithium acetate standard protocol. Before mating, yeast transformed with the WT interacting protein plasmid (**RPrey_S3**, MATα) was grown in non-selective medium for 12-18 h at 30°C, and concentrated to an OD_600_ of 40-80. Yeast containing mutant library DNA were collected in 1x nitrogen base (NB) media and mixed with yeast containing WT interacting protein at 2:1. The mixture was transferred to YPDA agar plates (six-well plate format) and incubated at 30°C. After 24 h, yeast was collected in 1x NB media, diluted and transferred to diploid non-selective NB-agar (on BioAssay 22×22 cm square dishes). After incubation at 30°C for 48 h, yeast was collected in 1x NB media, diluted to an OD_600_ of 0.2, and equally distributed on NB-agar containing amino acids and nucleic acids for reporter gene selection. Kinase expression was induced by addition of 200 μM copper sulfate (Merck KGaA, 7758-99-8) in the media. Following incubation at 30°C for 48-96 h, colonies were collected the plasmid DNA was isolated purified through phenol extraction and ethanol precipitation. PCR was performed with a proof-reading KOD polymerase (Sigma-Aldrich, 71086) using vector-specific primers.

### Sequence data analysis

Preparation (NextSeq High, 318 Cycles) for sequencing on an Illumina NextSeq 500 in a 150-base-pair paired-end read mode was done at sequencing core facility of the MPIMG, Berlin. Data analysis was performed using custom-made Perl and R scripts [44]. In brief, fastq files were converted to unique paired end sequence fasta files, followed by sequence alignment using STAR alignment software in paired-end mode [72]. Uniform distribution of mutations in the sequence and initial statistical analysis was evaluated. A linear model was used to calculate enrichment of individual mutations and a score was calculated for each

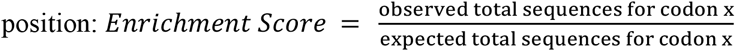

High confidence cut-offs of 50 sequences and enrichment above two-fold variation of the linear model were applied for coded mutations. In addition to the previously published code, refinements in the code were added. These included the following corrections: Reads with a high proportion of secondary mutations (> 0.8 of the maximal signal) present in combination with a given mutant were excluded, and insertions and deletions were locally removed from the sequences and confidence cutoffs implemented (< 0.7 of the maximal signal).

### Cell Culture

HEK293T (DSZM, ACC 872) cells were cultured in Dulbecco’s Modified Eagle’s Medium (DMEM, Thermo Fisher Scientific Inc., 41966-029) with 10% Fetal Bovine Serum (FBS, Thermo Fisher Scientific, 10270-106) and 1% Pen-Strep (10,000 U/ml, Thermo Fisher Scientific Inc., 15140-122). A549 (courtesy of ZMF cell line collection, MedUniGraz) cells were cultured in DMEM/F-12 Nutrient Mixture (1:1) (Thermo Fisher Scientific Inc., 31330-038) with 10% Fetal Bovine Serum (FBS, Thermo Fisher Scientific Inc., 10270-106) and 1% Pen-Strep (10,000 U/ml, Thermo Fisher Scientific Inc., 15140-122). Both cell lines were maintained in a humidified incubator at 37°C with 5% CO_2_.

### Cellular localisation experiments – Cell staining for fluorescence microscopy

24 h prior to transfection, HEK293T cells were seeded on poly-D-lysine (Sigma Aldrich, P7405) coated glass coverslips (15 mm, 24-well plate). The cells were transiently transfected with N-terminal YFP-fused NF2 variants (pdEYFP) and mutant proteins using polyethylenimine (PEI, 1 mg/ml, Alfa Aesar, 9002-98-6) at 1:5 ratio of pDNA:transfection reagent. 24 h afterwards, the cells were washed with PBS, pH 7.4 (Thermo Fisher Scientific Inc., 10010-015) and fixed with 4% paraformaldehyde (Sigma-Aldrich, 158127) solution (20 min at RT). Following three additional washing steps, cell membrane permeabilization was achieved by adding 0.1% Triton X-100 (Sigma-Aldrich, T8787) / 1 x PBS. The fixation agent was removed (three washing steps) and cells were counterstained with the following fluorescent dyes: Rhodamine-Phalloidin (Invitrogen, Thermo Fisher Scientific Inc., R415) and DAPI (4′,6-diamidino-2-phenylindole, Carl Roth, 6335) using a standard protocol. Following cell staining, the glass coverslips were mounted using Mounting medium (Sigma Aldrich, M1289), the rims were sealed with lacquer, and stored in the dark at 4°C until measurement. Images were taken with the following excitation (Ex) and emission (Em) settings: Rhodamine-Phalloidin: Ex 558 nm, Em 575-640 nm, DAPI: Ex 395 nm, Em 420-470 nm, EYFP signal: Ex 495 nm, Em 500-550 nm. All images were captured with a Zeiss AxioObserver Z1 microscope and a LD Plan-Neofluar 40x / 0.6 Korr Ph 2 objective combined with a Zeiss AxioCam MR. Image analysis was performed with the Zeiss ZEN 3.0 software package.

### Fluorescence-activated cell sorting (FACS) experiments

For FACS based proliferation analyses, 9 × 10^3^ HEK293T cells and 7-9 × 10^3^ A549 cells were plated in each well of a 96-well plate and incubated for 24 h prior to transfection. The cells were transiently transfected with 150 ng plasmid DNA of YFP-fused NF2 WT and mutant constructs using PEI (1 mg/ml) at a ratio of 1:5 and 1:4, respectively. Individual experiments were carried out in triplicates. Complete growth medium was exchanged 12- and 48 h post-transfection. Cells were trypsinized (Trypsin-EDTA, 0.25% (Thermo Fisher Scientific Inc., 25200-056) and 0.5% (Thermo Fisher Scientific Inc., 15400-054), respectively) according to a standard protocol and resuspended in 70-100 μl (according to cell density) ice-cold 1 mM FACS buffer for HEK293T and 2.5 mM FACS buffer for A549 cells (FACS buffer: 1-2.5 mM EDTA, pH 8.0 and 2% FBS in 1 x PBS). In four experiments, the relative fraction of transfected YFP-positive cells was recorded every 24 h over a period of three days. Before starting the flow cytometry measurements performed on a BD FACS Fortessa or a Cytek Aurora flow cytometer, the cells were stained with 1 μl of propidium iodide (PI (MP Biomedicals, LLC, 195458), 1 mg/ml in DMSO (Sigma-Aldrich, D8418)) to distinguish dead cells. The percentage of YFP-positive cells was normalized across the fifteen variants and expressed as Z-score. A cutoff of -0.8 or 0.8 was set. The following laser and filter wavelengths were used for the flow cytometry measurements; PI fluorophore: 561 nm, 670/30 nm bandpass filter; and YFP fluorophore: 488 nm, 510/20 nm bandpass filter. The flow rate was adjusted to <1000 events/second.

### IncuCyte live cell proliferation assays

HEK293T, A549 and SC4 cells were cultured in DMEM supplemented with 10% FBS at 37°C and 5% CO_2_ atmosphere. Lipofections were performed using jetPRIME (Polyplus, VWR 89129-920) or Lipofectamine 3000 (ThermoFisher, L3000001) reagent according to the manufacturer’s instructions. Cells were seeded in flat-bottom, transparent 96-well plates at approximately 3-4 × 10^3^ cells per well. 24 h post seeding, the HEK293T cells were transfected with 50 ng/well and the SC4 cells with 100 ng/well of the respective DNA constructs. To reduce cytotoxicity, the medium was changed 4 h post transfection. Starting at 24 h post transfection, live-cell proliferation and expression of YFP-tagged proteins were tracked for 48 h using an Incucyte S3 device (Sartorius).

## Data accessibility

Raw data are accessible via ENA Project Accession Number PRJEB57973 at https://www.ebi.ac.uk/ena/browser/home.

## Author contributions

Conceptualization: US

Data curation: CSM, JW

Formal Analysis: JW, CSM

Funding acquisition: US, ES

Investigation: CSM, JW, EA, AF, EJ-L, SJ, JMW, CLH

Methodology: CSM, JW, ES, US

Project administration:

Resources: ES, US

Software: JW

Supervision: ES, US

Validation: CSM, EA, AF, EJ-L

Visualization: CSM, JW, US

Writing – original draft: US, CSM

Writing – review & editing: US, CSM and all authors

## Acknowledgements

We thank Nouhad Benlasfer and Natalia Kunowska for help with the experiments. We thank Bernd Timmermann and the members of MPI-MG Sequencing Facility for performing the second generation sequencing experiments. SC4 cells were obtained as courtesy from Helen Morrison (Leibniz Institute on Aging, Jena). NF2 isoform cDNAs were obtained as courtesy of Michael Kressel (University of Erlangen). The work was funded by the Austrian Science Fund (FWF, project P30162). The work was supported by the FWF doc.fund Molecular Metabolism (DOC 50) and the Field of Excellence BioHealth -University of Graz. ES was supported by grants from the FWF P30441, P32960, P35159 and the Tyrolean Cancer Society.

## Figure Legends

**Supplemental Figure S1:**
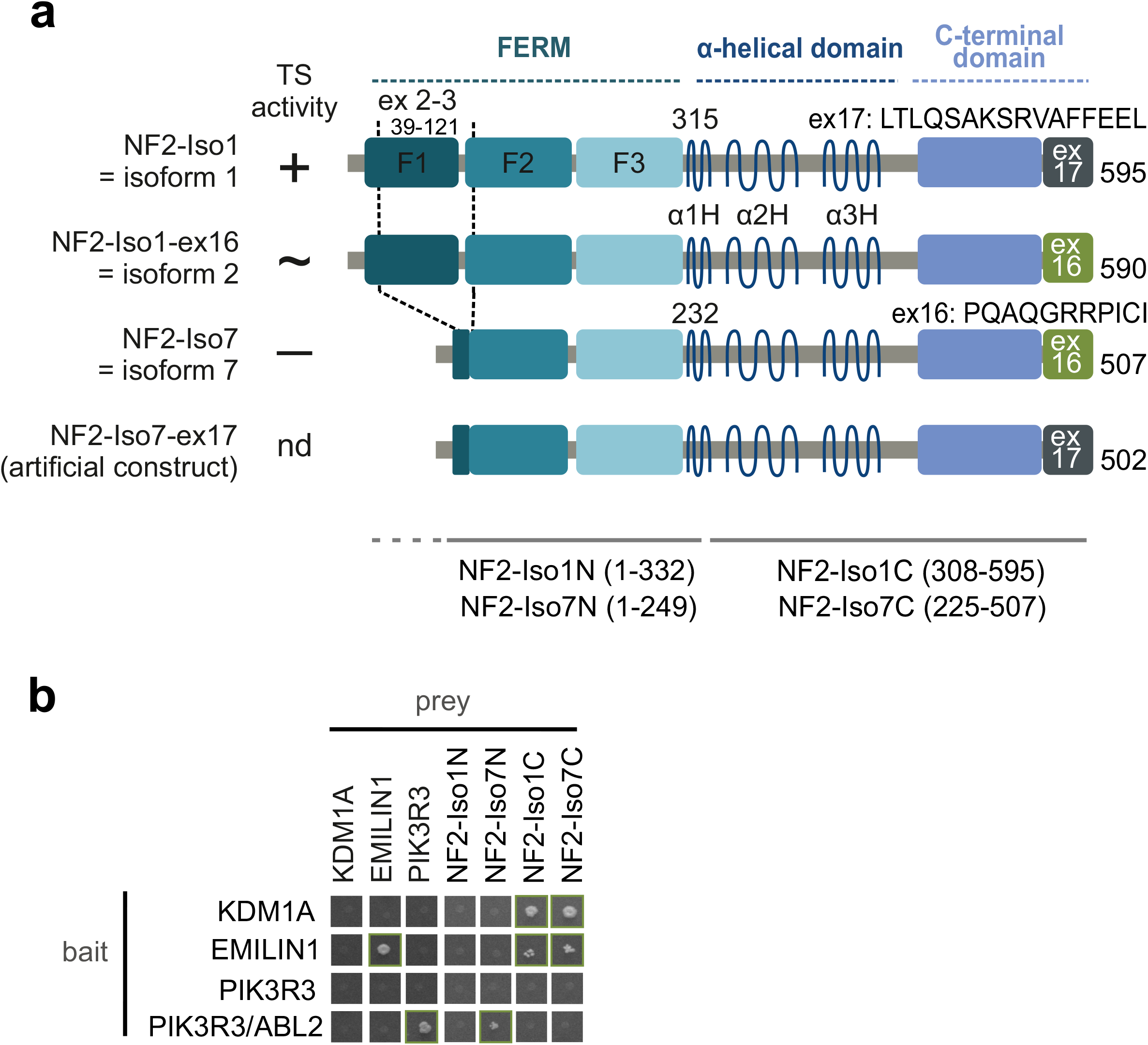
**a**. NF2 splice isoforms and constructs used in the study. The canonical isoform 1 is 595 AA long and contains exon 17 (P35240-1). It includes all features of isoforms 2 and 7 except for differences in the C-terminal end from amino acids 580-595. Isoform 2 (P35240-2) has a length of 590 amino acids and differs in the C-terminal end derived by alternative splicing of exon 16 instead of exon 17 from isoform 1. Isoform 7 is a shorter splice isoform with 507 amino acids, lacking the majority of the F1 FERM subdomain and the beginning of the F2 subdomain (exons 2 and 3, AA 39-121, P35240-4). As for isoform 2, the C-terminal end consists of exon 16. Several other splice isoforms were described, in addition to the three isoforms depicted, however NF2-Iso7-ex17 is an artificial construct to test the influence of the isoform 1 C-terminus on the interactions with the shorter isoform 7. Constructs representing the N-terminal and C-terminal halves of NF2 are indicated. Tumor suppressor activity (TS activity) was demonstrated for isoform 1 (+) but not for isoform 7 (-); nd = not determined. **b**. Y2H protein interaction results. Y2H mapping of NF2 interacting regions. Growth on selective agar, indicating protein interaction is shown. EMILIN1 formed homo dimers and PIK3R3 formed a homodimeric interaction in the presence of an active tyrosine kinase ABL2. This requirement for a human tyrosine kinase explains the requirement for ABL2 for the interaction of PIK3R3 with wild-type NF2-Iso7N, the N-terminal half of isoform 7. The reverse bait-prey configuration of the experiment is shown in Figure 1. KDM1A and EMILIN1 both interact with the NF2 C-terminal halves, independently of whether the last amino acids resemble isoform 1 or isoform 7. Please note that NF2-Iso1C and NF2-Iso7C were autoactive when used as bait and were therefore excluded from the analysis.

**Supplemental Figure S2:**
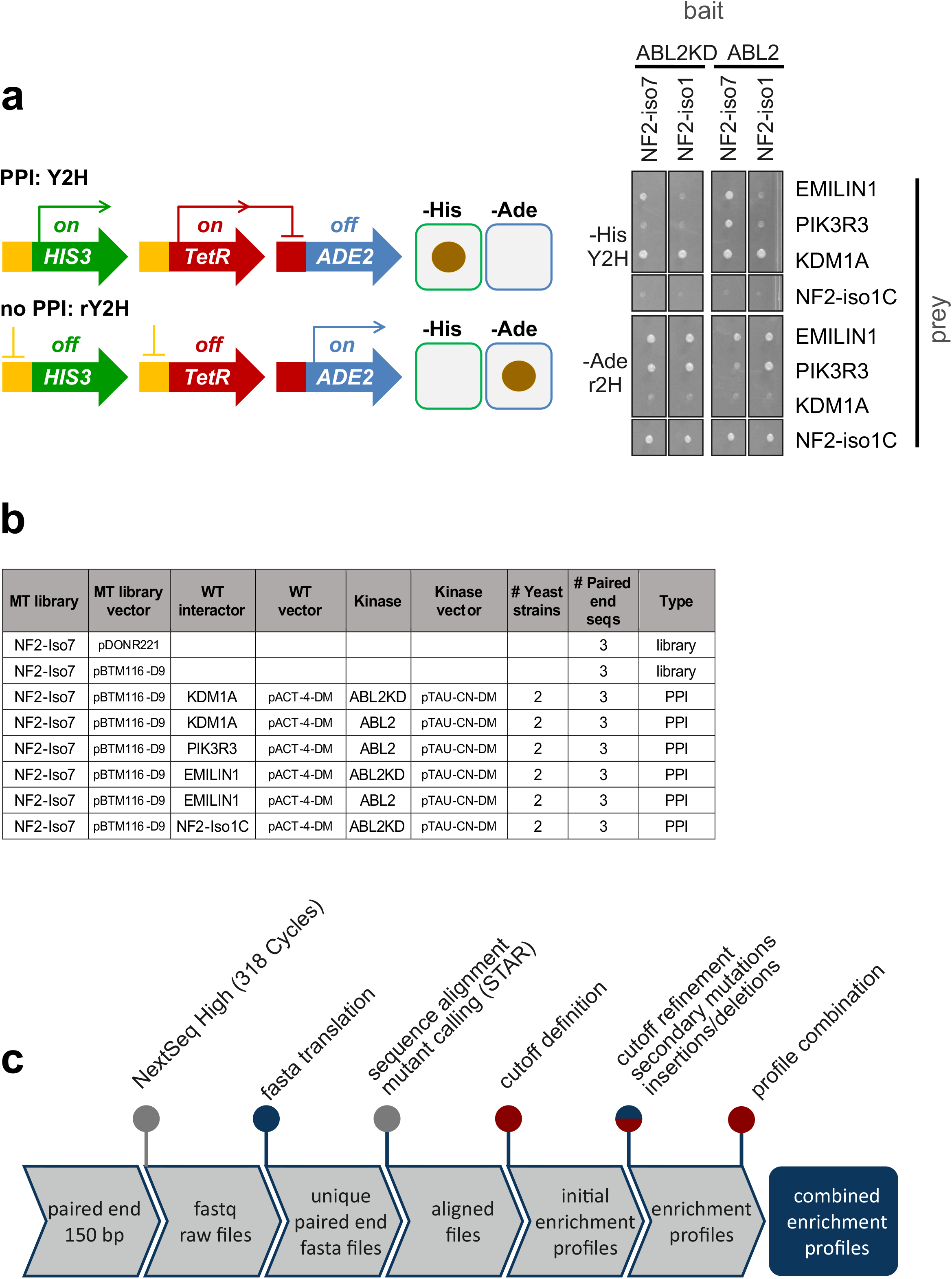
**a**. Reverse Y2H analysis. Genetics of reverse-Y2H strain (left). A bait-prey pair can be assayed for interactions on selective agar lacking histidine or lacking adenine respectively. Interacting bait-prey pairs grow on agar lacking histidine (-his) and non-interacting pairs grown on agar lacking adenine (-ade). When using a deep mutational scanning pool representing all single amino acid exchanges in the place of one partner, non-interacting variants can be selected on agar lacking adenine. A representative growth experiment (right) growing the strains on the two agar plates, lacking adenine (rY2H) or histidine (Y2H) respectively. **b**. Overview of the rY2H deep mutagenesis interaction experiments and the sequencing runs used for evaluation. All MT libraries generated were sequenced three times in the entry vector and in the yeast expression vector as controls. The controls did not face selection pressure and the combined sequence read count was used as baseline for the enrichment analysis of the rY2H interactions. The rY2H interactions were tested in two yeast strain pairs and three biological replicates were individually sequenced resulting in a total of six sequencing runs per interaction. **c**. Sequence analysis pipeline for the generation of mutant enrichment profiles. Blue dots represent Perl scripts, red dots indicate R script used in the analysis step. Sequence alignment and mutant calling was done with the STAR package.

**Supplemental Figure S3:**
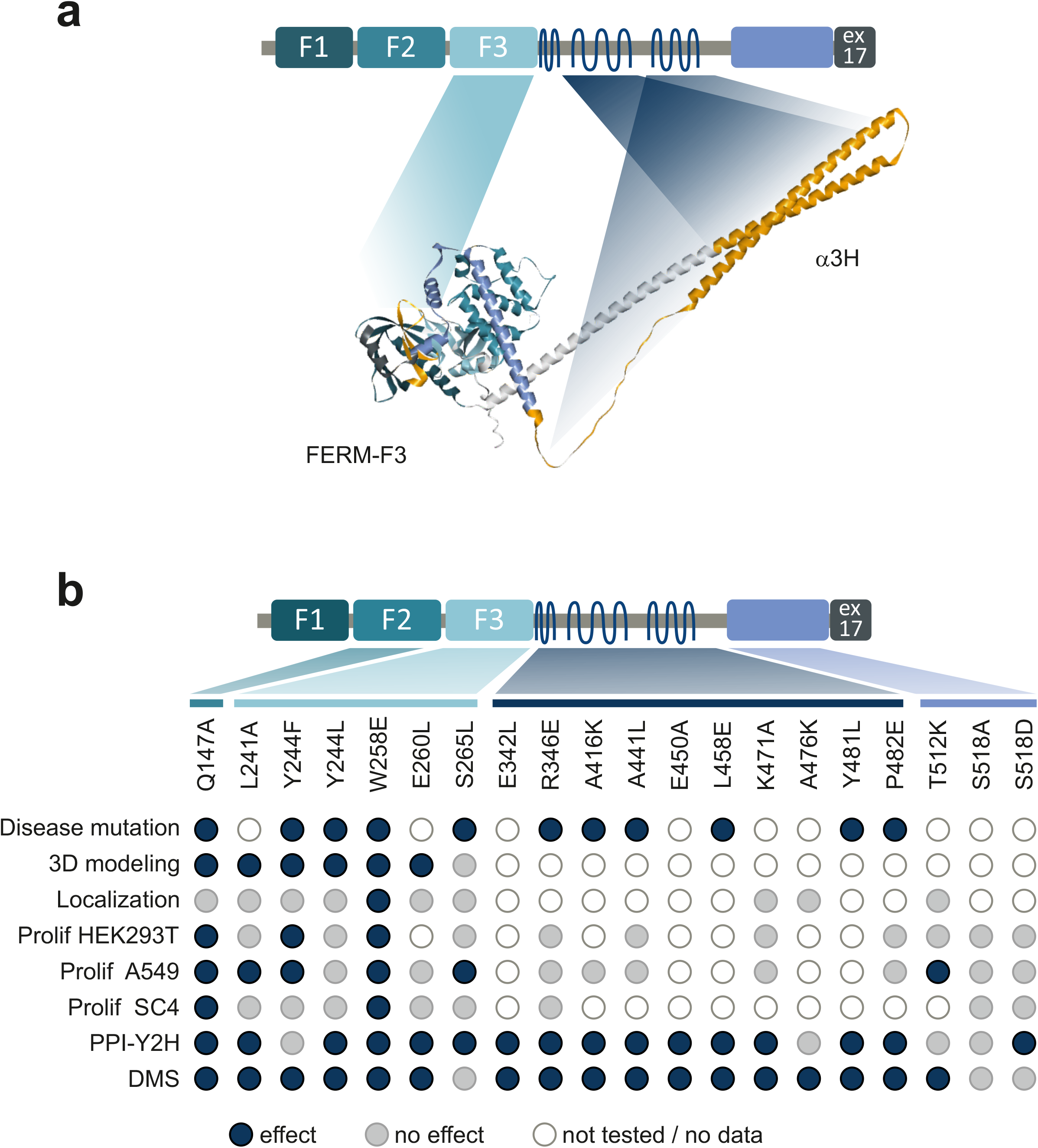
**a**. Schematic and 3D model of NF2 summarizing of regions critical for NF2 conformation. FERM-F3 and the α2H / α3H helix regions harboring mutations critical for conformation dependent interactions are colored in orange in the alpha fold 3D structure (AF P35240 F1). **b**. Summary of the NF2 variant effects. Disease mutations: a blue dot indicates an annotated SUV at the amino acid position. 3D modelling: a blue dot indicates a change of 3D interactions for the amino acid substitution. Localization: blue indicates a changed subcellular distribution for the amino acid substitution. Prolif: a blue dot indicates altered proliferation upon NF2 variant expression of HEK293T, A459 or SC4 cells, respectively. PPI-Y2H: a blue dot indicates alter protein interaction patterns in Y2H spot assays with individually cloned NF2 variants. DMS: a blue dot indicates interaction perturbation of the amino acid substitution in deep scanning experiments. Grey, no effect in comparison to wild-type; White / no fill, not tested.

**Supplemental Table S1**

Interaction perturbation rY2H scores of the NF2 isoform 7 with its partners

## References

1. Bretscher A, Edwards K, Fehon RG. ERM proteins and merlin: integrators at the cell cortex. Nat Rev Mol Cell Biol. 2002;3:586–99. doi:10.1038/nrm882.

2. Laulajainen M, Muranen T, Carpén O, Grönholm M. Protein kinase A-mediated phosphorylation of the NF2 tumor suppressor protein merlin at serine 10 affects the actin cytoskeleton. Oncogene. 2008;27:3233–43. doi:10.1038/sj.onc.1210988.

3. Rouleau GA, Merel P, Lutchman M, Sanson M, Zucman J, Marineau C, et al. Alteration in a new gene encoding a putative membrane-organizing protein causes neuro-fibromatosis type 2. Nature. 1993;363:515–21. doi:10.1038/363515a0.

4. Trofatter J. A novel moesin-, ezrin-, radixin-like gene is a candidate for the neurofibromatosis 2 tumor suppressor. Cell. 1993;72:791–800. doi:10.1016/0092-8674(93)90406-g.

5. Morrison H, Sherman LS, Legg J, Banine F, Isacke C, Haipek CA, et al. The NF2 tumor suppressor gene product, merlin, mediates contact inhibition of growth through interactions with CD44. Genes Dev. 2001;15:968–80. doi:10.1101/gad.189601.

6. Okada T, Lopez-Lago M, Giancotti FG. Merlin/NF-2 mediates contact inhibition of growth by suppressing recruitment of Rac to the plasma membrane. J Cell Biol. 2005;171:361–71. doi:10.1083/jcb.200503165.

7. Curto M, Cole BK, Lallemand D, Liu C-H, McClatchey AI. Contact-dependent inhibition of EGFR signaling by Nf2/Merlin. J Cell Biol. 2007;177:893–903. doi:10.1083/jcb.200703010.

8. Okada M, Wang Y, Jang S-W, Tang X, Neri LM, Ye K. Akt phosphorylation of merlin enhances its binding to phosphatidylinositols and inhibits the tumor-suppressive activities of merlin. Cancer Res. 2009;69:4043–51. doi:10.1158/0008-5472.CAN-08-3931.

9. Lallemand D, Curto M, Saotome I, Giovannini M, McClatchey AI. NF2 deficiency promotes tumorigenesis and metastasis by destabilizing adherens junctions. Genes Dev. 2003;17:1090–100. doi:10.1101/gad.1054603.

10. Lallemand D, Manent J, Couvelard A, Watilliaux A, Siena M, Chareyre F, et al. Merlin regulates transmembrane receptor accumulation and signaling at the plasma membrane in primary mouse Schwann cells and in human schwannomas. Oncogene. 2009;28:854–65. doi:10.1038/onc.2008.427.

11. Morrison H, Sperka T, Manent J, Giovannini M, Ponta H, Herrlich P. Merlin/neurofibromatosis type 2 suppresses growth by inhibiting the activation of Ras and Rac. Cancer Res. 2007;67:520–7. doi:10.1158/0008-5472.CAN-06-1608.

12. Cui Y, Groth S, Troutman S, Carlstedt A, Sperka T, Riecken LB, et al. The NF2 tumor suppressor merlin interacts with Ras and RasGAP, which may modulate Ras signaling. Oncogene. 2019;38:6370–81. doi:10.1038/s41388-019-0883-6.

13. Hamaratoglu F, Willecke M, Kango-Singh M, Nolo R, Hyun E, Tao C, et al. The tumour-suppressor genes NF2/Merlin and Expanded act through Hippo signalling to regulate cell proliferation and apoptosis. Nat Cell Biol. 2006;8:27–36. doi:10.1038/ncb1339.

14. Li W, You L, Cooper J, Schiavon G, Pepe-Caprio A, Zhou L, et al. Merlin/NF2 suppresses tumorigenesis by inhibiting the E3 ubiquitin ligase CRL4(DCAF1) in the nucleus. Cell. 2010;140:477–90. doi:10.1016/j.cell.2010.01.029.

15. Zhang N, Bai H, David KK, Dong J, Zheng Y, Cai J, et al. The Merlin/NF2 tumor suppressor functions through the YAP oncoprotein to regulate tissue homeostasis in mammals. Dev Cell. 2010;19:27–38. doi:10.1016/j.devcel.2010.06.015.

16. Cooper J, Giancotti FG. Molecular insights into NF2/Merlin tumor suppressor function. FEBS Lett. 2014;588:2743–52. doi:10.1016/j.febslet.2014.04.001.

17. Meng F, Yu Z, Zhang D, Chen S, Guan H, Zhou R, et al. Induced phase separation of mutant NF2 imprisons the cGAS-STING machinery to abrogate antitumor immunity. Mol Cell. 2021;81:4147-4164.e7. doi:10.1016/j.molcel.2021.07.040.

18. Asthagiri AR, Parry DM, Butman JA, Kim HJ, Tsilou ET, Zhuang Z, Lonser RR. Neurofibromatosis type 2. The Lancet. 2009;373:1974–86. doi:10.1016/S0140-6736(09)60259-2.

19. Evans DGR. Neurofibromatosis type 2 (NF2): a clinical and molecular review. Orphanet J Rare Dis. 2009;4:16. doi:10.1186/1750-1172-4-16.

20. Petrilli AM, Fernández-Valle C. Role of Merlin/NF2 inactivation in tumor biology. Oncogene. 2016;35:537–48. doi:10.1038/onc.2015.125.

21. Martincorena I, Raine KM, Gerstung M, Dawson KJ, Haase K, van Loo P, et al. Universal Patterns of Selection in Cancer and Somatic Tissues. Cell. 2017;171:1029-1041.e21. doi:10.1016/j.cell.2017.09.042.

22. Vogelstein B, Papadopoulos N, Velculescu VE, Zhou S, Diaz LA, Kinzler KW. Cancer genome landscapes. Science. 2013;339:1546–58. doi:10.1126/science.1235122.

23. Sherman L, Xu HM, Geist RT, Saporito-Irwin S, Howells N, Ponta H, et al. Interdomain binding mediates tumor growth suppression by the NF2 gene product. Oncogene. 1997;15:2505–9. doi:10.1038/sj.onc.1201418.

24. Scoles DR, Chen M, Pulst S-M. Effects of Nf2 missense mutations on schwannomin interactions. Biochem Biophys Res Commun. 2002;290:366–74. doi:10.1006/bbrc.2001.6178.

25. Sher I, Hanemann CO, Karplus PA, Bretscher A. The tumor suppressor merlin controls growth in its open state, and phosphorylation converts it to a less-active more-closed state. Dev Cell. 2012;22:703–5. doi:10.1016/j.devcel.2012.03.008.

26. Deguen B, Mérel P, Goutebroze L, Giovannini M, Reggio H, Arpin M, Thomas G. Impaired interaction of naturally occurring mutant NF2 protein with actin-based cytoskeleton and membrane. Hum Mol Genet. 1998;7:217–26. doi:10.1093/hmg/7.2.217.

27. Koga H, Araki N, Takeshima H, Nishi T, Hirota T, Kimura Y, et al. Impairment of cell adhesion by expression of the mutant neurofibromatosis type 2 (NF2) genes which lack exons in the ERM-homology domain. Oncogene. 1998;17:801–10. doi:10.1038/sj.onc.1202010.

28. Giovannini M, Robanus-Maandag E, Niwa-Kawakita M, van der Valk M, Woodruff JM, Goutebroze L, et al. Schwann cell hyperplasia and tumors in transgenic mice expressing a naturally occurring mutant NF2 protein. Genes Dev. 1999;13:978–86. doi:10.1101/gad.13.8.978.

29. LaJeunesse DR, McCartney BM, Fehon RG. Structural analysis of Drosophila merlin reveals functional domains important for growth control and subcellular localization. J Cell Biol. 1998;141:1589–99. doi:10.1083/jcb.141.7.1589.

30. Johnson KC, Kissil JL, Fry JL, Jacks T. Cellular transformation by a FERM domain mutant of the Nf2 tumor suppressor gene. Oncogene. 2002;21:5990–7. doi:10.1038/sj.onc.1205693.

31. Luo Z-L, Cheng S-Q, Shi J, Zhang H-L, Zhang C-Z, Chen H-Y, et al. A splicing variant of Merlin promotes metastasis in hepatocellular carcinoma. Nat Commun. 2015;6:8457. doi:10.1038/ncomms9457.

32. Grönholm M, Sainio M, Zhao F, Heiska L, Vaheri A, Carpén O. Homotypic and heterotypic interaction of the neurofibromatosis 2 tumor suppressor protein merlin and the ERM protein ezrin. J Cell Sci. 1999;112 (Pt 6):895–904. doi:10.1242/jcs.112.6.895.

33. Gutmann DH, Haipek CA, Hoang Lu K. Neurofibromatosis 2 tumor suppressor protein, merlin, forms two functionally important intramolecular associations. J Neurosci Res. 1999;58:706–16.

34. Brault E, Gautreau A, Lamarine M, Callebaut I, Thomas G, Goutebroze L. Normal membrane localization and actin association of the NF2 tumor suppressor protein are dependent on folding of its N-terminal domain. J Cell Sci. 2001;114:1901–12. doi:10.1242/jcs.114.10.1901.

35. Kissil JL, Johnson KC, Eckman MS, Jacks T. Merlin phosphorylation by p21-activated kinase 2 and effects of phosphorylation on merlin localization. J Biol Chem. 2002;277:10394–9. doi:10.1074/jbc.M200083200.

36. Xiao G-H, Beeser A, Chernoff J, Testa JR. p21-activated kinase links Rac/Cdc42 signaling to merlin. J Biol Chem. 2002;277:883–6. doi:10.1074/jbc.C100553200.

37. Shaw RJ, Paez JG, Curto M, Yaktine A, Pruitt WM, Saotome I, et al. The Nf2 Tumor Suppressor, Merlin, Functions in Rac-Dependent Signaling. Dev Cell. 2001;1:63–72. doi:10.1016/s1534-5807(01)00009-0.

38. Hennigan RF, Foster LA, Chaiken MF, Mani T, Gomes MM, Herr AB, Ip W. Fluorescence resonance energy transfer analysis of merlin conformational changes. Mol Cell Biol. 2010;30:54–67. doi:10.1128/MCB.00248-09.

39. Meng J-J, Lowrie DJ, Sun H, Dorsey E, Pelton PD, Bashour A-M, et al. Interaction between two isoforms of the NF2 tumor suppressor protein, merlin, and between merlin and ezrin, suggests modulation of ERM proteins by merlin. J Neurosci Res. 2000;62:491–502. doi:10.1002/1097-4547(20001115)62:4<491::AID-JNR3>3.0.CO;2-D.

40. Phang JM, Harrop SJ, Duff AP, Sokolova AV, Crossett B, Walsh JC, et al. Structural characterization suggests models for monomeric and dimeric forms of full-length ezrin. Biochem J. 2016;473:2763–82. doi:10.1042/BCJ20160541.

41. Esposito D, Weile J, Shendure J, Starita LM, Papenfuss AT, Roth FP, et al. MaveDB: an open-source platform to distribute and interpret data from multiplexed assays of variant effect. Genome Biol. 2019;20:223. doi:10.1186/s13059-019-1845-6.

42. Woodsmith J, Stelzl U. Understanding Disease Variants through the Lens of Protein Interactions. Cell Syst. 2017;5:544–6. doi:10.1016/j.cels.2017.12.009.

43. Yadav A, Vidal M, Luck K. Precision medicine - networks to the rescue. Curr Opin Biotechnol. 2020;63:177–89. doi:10.1016/j.copbio.2020.02.005.

44. Woodsmith J, Apelt L, Casado-Medrano V, Özkan Z, Timmermann B, Stelzl U. Protein interaction perturbation profiling at amino-acid resolution. Nat Methods. 2017;14:1213–21. doi:10.1038/nmeth.4464.

45. Grossmann A, Benlasfer N, Birth P, Hegele A, Wachsmuth F, Apelt L, Stelzl U. Phospho-tyrosine dependent protein-protein interaction network. Mol Syst Biol. 2015;11:794. doi:10.15252/msb.20145968.

46. Jehle S, Kunowska N, Benlasfer N, Woodsmith J, Weber G, Wahl MC, Stelzl U. A human kinase yeast array for the identification of kinases modulating phosphorylation-dependent protein-protein interactions; 2021.

47. Weimann M, Grossmann A, Woodsmith J, Özkan Z, Birth P, Meierhofer D, et al. A Y2H-seq approach defines the human protein methyltransferase interactome. Nat Methods. 2013;10:339–42. doi:10.1038/nmeth.2397.

48. Haenig C, Atias N, Taylor AK, Mazza A, Schaefer MH, Russ J, et al. Interactome Mapping Provides a Network of Neurodegenerative Disease Proteins and Uncovers Widespread Protein Aggregation in Affected Brains. Cell Rep. 2020;32:108050. doi:10.1016/j.celrep.2020.108050.

49. Go CD, Knight JDR, Rajasekharan A, Rathod B, Hesketh GG, Abe KT, et al. A proximity-dependent biotinylation map of a human cell. Nature. 2021;595:120–4. doi:10.1038/s41586-021-03592-2.

50. Kitzman JO, Starita LM, Lo RS, Fields S, Shendure J. Massively parallel single-amino-acid mutagenesis. Nat Methods. 2015;12:203-6, 4 p following 206. doi:10.1038/nmeth.3223.

51. Li Y, Zhou H, Li F, Chan SW, Lin Z, Wei Z, et al. Angiomotin binding-induced activation of Merlin/NF2 in the Hippo pathway. Cell Res. 2015;25:801–17. doi:10.1038/cr.2015.69.

52. Chinthalapudi K, Mandati V, Zheng J, Sharff AJ, Bricogne G, Griffin PR, et al. Lipid binding promotes the open conformation and tumor-suppressive activity of neurofibromin 2. Nat Commun. 2018;9:1338. doi:10.1038/s41467-018-03648-4.

53. Primi MC, Rangarajan ES, Patil DN, Izard T. Conformational flexibility determines the Nf2/merlin tumor suppressor functions. Matrix Biol Plus. 2021;12:100074. doi:10.1016/j.mbplus.2021.100074.

54. Xing W, Li M, Zhang F, Ma X, Long J, Zhou H. The conformation change and tumor suppressor role of Merlin are both independent of Serine 518 phosphorylation. Biochem Biophys Res Commun. 2017;493:46–51. doi:10.1016/j.bbrc.2017.09.077.

55. Michie KA, Bermeister A, Robertson NO, Goodchild SC, Curmi PMG. Two Sides of the Coin: Ezrin/Radixin/Moesin and Merlin Control Membrane Structure and Contact Inhibition. Int J Mol Sci 2019. doi:10.3390/ijms20081996.

56. Surace EI, Haipek CA, Gutmann DH. Effect of merlin phosphorylation on neurofibromatosis 2 (NF2) gene function. Oncogene. 2004;23:580–7. doi:10.1038/sj.onc.1207142.

57. Ali Khajeh J, Ju JH, Atchiba M, Allaire M, Stanley C, Heller WT, et al. Molecular conformation of the full-length tumor suppressor NF2/Merlin--a small-angle neutron scattering study. J Mol Biol. 2014;426:2755–68. doi:10.1016/j.jmb.2014.05.011.

58. Kang BS, Cooper DR, Devedjiev Y, Derewenda U, Derewenda ZS. The structure of the FERM domain of merlin, the neurofibromatosis type 2 gene product. Acta Crystallogr D Biol Crystallogr. 2002;58:381–91. doi:10.1107/s0907444901021175.

59. Li Y, Wei Z, Zhang J, Yang Z, Zhang M. Structural basis of the binding of Merlin FERM domain to the E3 ubiquitin ligase substrate adaptor DCAF1. J Biol Chem. 2014;289:14674–81. doi:10.1074/jbc.M114.551184.

60. Mori T, Gotoh S, Shirakawa M, Hakoshima T. Structural basis of DDB1-and-Cullin 4-associated Factor 1 (DCAF1) recognition by merlin/NF2 and its implication in tumorigenesis by CD44-mediated inhibition of merlin suppression of DCAF1 function. Genes Cells. 2014;19:603–19. doi:10.1111/gtc.12161.

61. Nguyen R, Reczek D, Bretscher A. Hierarchy of merlin and ezrin N- and C-terminal domain interactions in homo- and heterotypic associations and their relationship to binding of scaffolding proteins EBP50 and E3KARP. J Biol Chem. 2001;276:7621–9. doi:10.1074/jbc.M006708200.

62. Kunowska N, Stelzl U. Decoding the cellular effects of genetic variation through interaction proteomics. Curr Opin Chem Biol. 2021;66:102100. doi:10.1016/j.cbpa.2021.102100.

63. Starita LM, Ahituv N, Dunham MJ, Kitzman JO, Roth FP, Seelig G, et al. Variant Interpretation: Functional Assays to the Rescue. Am J Hum Genet. 2017;101:315–25. doi:10.1016/j.ajhg.2017.07.014.

64. Starita LM, Young DL, Islam M, Kitzman JO, Gullingsrud J, Hause RJ, et al. Massively Parallel Functional Analysis of BRCA1 RING Domain Variants. Genetics. 2015;200:413–22. doi:10.1534/genetics.115.175802.

65. Faure AJ, Domingo J, Schmiedel JM, Hidalgo-Carcedo C, Diss G, Lehner B. Mapping the energetic and allosteric landscapes of protein binding domains. Nature. 2022;604:175–83. doi:10.1038/s41586-022-04586-4.

66. Bolognesi B, Faure AJ, Seuma M, Schmiedel JM, Tartaglia GG, Lehner B. The mutational landscape of a prion-like domain. Nat Commun. 2019;10:4162. doi:10.1038/s41467-019-12101-z.

67. Findlay GM, Daza RM, Martin B, Zhang MD, Leith AP, Gasperini M, et al. Accurate classification of BRCA1 variants with saturation genome editing. Nature. 2018;562:217–22. doi:10.1038/s41586-018-0461-z.

68. Matreyek KA, Starita LM, Stephany JJ, Martin B, Chiasson MA, Gray VE, et al. Multiplex assessment of protein variant abundance by massively parallel sequencing. Nat Genet. 2018;50:874–82. doi:10.1038/s41588-018-0122-z.

69. Cole BK, Curto M, Chan AW, McClatchey AI. Localization to the cortical cytoskeleton is necessary for Nf2/merlin-dependent epidermal growth factor receptor silencing. Mol Cell Biol. 2008;28:1274–84. doi:10.1128/MCB.01139-07.

70. Mani T, Hennigan RF, Foster LA, Conrady DG, Herr AB, Ip W. FERM domain phosphoinositide binding targets merlin to the membrane and is essential for its growth-suppressive function. Mol Cell Biol. 2011;31:1983–96. doi:10.1128/MCB.00609-10.

71. Landrum MJ, Chitipiralla S, Brown GR, Chen C, Gu B, Hart J, et al. ClinVar: improvements to accessing data. Nucleic Acids Res. 2020;48:D835–D844. doi:10.1093/nar/gkz972.

72. Dobin A, Davis CA, Schlesinger F, Drenkow J, Zaleski C, Jha S, et al. STAR: ultrafast universal RNA-seq aligner. Bioinformatics. 2013;29:15–21. doi:10.1093/bioinformatics/bts635.

